# Microbial Metamorphosis: Symbiotic bacteria and fungi proliferate during diapause and may enhance overwintering survival in a solitary bee

**DOI:** 10.1101/2023.11.02.565352

**Authors:** S. M. Christensen, S. Srinivas, Q. McFrederick, B. Danforth, S. L. Buchmann, R. L. Vannette

## Abstract

Host-microbe interactions underlie the development and fitness of many macroorganisms including bees. While many social bees benefit from vertically transmitted gut bacteria, solitary bees, which comprise the vast majority of species diversity within bees, lack a specialized gut microbiome. Here we examine the composition and abundance of bacteria and fungi throughout the complete life cycle of a ground-nesting solitary bee *Anthophora bomboides standfordiana*. In contrast to expectations, immature bee stages maintain a distinct core microbiome consisting of Actinobacteria and fungi in the genus *Moniliella*. Diapausing larval bees hosted the most abundant and distinctive bacteria and fungi, attaining 33 and 52 times their initial copy number, respectively. We tested two adaptive hypotheses regarding microbial functions for overwintering bees. First, using isolated bacteria and fungi, we found that *Streptomyces* from brood cells inhibited the growth of multiple pathogenic filamentous fungi, suggesting a role in pathogen protection during the long period of diapause. Second, sugar alcohol composition changed in tandem with major changes in microbial abundance suggesting links with bee metabolism or overwintering biology. Our results suggest that *Anthophora bomboides* hosts a conserved core microbiome that may provide key fitness advantages through larval development and overwintering, and raises the question of how this microbiome is transmitted between generations. The present work suggests that focus on adult gut microbiomes in solitary bees may overlook microbial symbionts within brood cells that could play diverse roles in bee fitness, and that exploration of microbes associated with immature bees may uncover novel microbial effects on insect hosts.

## Introduction

The success of a wide range of insect species has hinged on partnerships with microbes (1, 2). The unique metabolic abilities of bacteria and fungi can facilitate novel resource use for insect hosts use *via* synthesis of limiting nutrients (3), evasion or detoxification of diet defenses (4) or in some cases cultivated as food themselves (5). Bacteria and fungi may also partner with insects in defensive symbiosis, providing defense from predation (6, 7), pathogens (8, 9), or food spoilage (10) through production of antagonistic or deterrent compounds, acidification or direct competition with insect pathogens.

Social corbiculate bees (e.g. honey bees, stingless bees, bumblebees), host well-characterized gut bacterial microbiomes which assist in the breakdown and detoxification of pollen, provision of essential steroids, regulation of immunity, and protection from pathogens (11–13). As in most host-microbe partnerships, vertical or social transmission is considered a key facilitator of this relationship by ensuring continuity of microbial associations between generations and thereby allowing for coevolution of host and microbe (12). Although the microbiomes of social bees have received the most study, the vast majority of bee species are solitary and do not engage in cooperative brood care nor shared foraging and feeding (14). Moreover, the solitary bee species studied to date generally lack consistent species-specific microbiomes (15, 16) in stark contrast to the conserved gut microbial communities found social bee species. While in the brood cells of some bee species, stored nectar and pollen can be dominated by lactobacilli these bacteria do not persist or are detected in very low abundance in the bee gut after defecation and during larval diapause (17–19). The combination of an annual phenology, solitary nesting, holometabolous development (complete metamorphosis), and lack of direct brood care may impede the persistence and transmission of a specialized core microbiome (20, 21). Instead, solitary bees are thought to acquire and filter microbes from the environment anew each generation (excepting occasional endosymbiotic bacteria, see (22)), resulting in variable microbial communities among individuals and populations.

The average solitary bee spends as much as 80% of its life developing inside a sealed brood cell, which is most often in an underground nest dug by the adult female bee (14). Brood cells are provisioned by the adult female with all of the pollen and nectar the developing bee will consume, then the egg is laid on the food provisions and the cell is sealed. Larvae consume the nectar and pollen, defecate, then pupate, and emerge the following year as adults (14). During this time, the developing bee undergoes significant changes in metabolism, morphology and likely immune function (14, 23–25). Notably, the developing bee is vulnerable to mold/filamentous fungi-which are estimated to cause significant mortality, ranging from 20% to 70% in ground-nesting bee species (26). Fungal brood pathogens such as chalk brood (*Ascosphaera*) thrive in cool, wet environments, as moisture promotes fungal growth and cooler temperatures can stress larvae, making them more vulnerable to infection (27, 28). Many bees line brood cells with linings originating from Dufour’s gland secretions, which are believed to have antimicrobial properties (14). Other ground nesting hymenopterans, including ants and wasps, are known to associate with Actinobacteria that produce antifungal compounds (8, 9, 29). However, to our knowledge such defensive mutualism has not been documented in solitary bee species.

Here, we characterize the composition, abundance, and potential functions of the bacterial and fungal microbiome of the solitary bee *Anthophora bomboides stanfordiana* Kirby, 1837 (Hymenoptera: Apidae) over 8 developmental stages, from two geographic sites, over two years using amplicon sequencing, qPCR, microbial isolations, and *in vitro* trials and assays. We describe a uniquely consistent core microbiome associated throughout development of this solitary bee species, how it changes through bee developmental stages, and finally test adaptive hypotheses regarding microbial effects on bee ecology and metabolism.

### Study system: bee biology and life history

*Anthophora bomboides stanfordiana* (from here: *A. bomboides)* is a gregariously nesting solitary bee, inhabiting coastal and inland bluffs along the western coast of North America. Adult females are mimics of the native *Bombus vosnesenskii*, and nest in densely populated sites containing tens to over a hundred thousand bees (30).

Nests are dug anew each year by females in the spring (Fig. 1). Female bees gather fresh water from a nearby seep or creek and wet the hard dirt to facilitate digging (27). A turret is constructed over the nest entrance with the freshly dug soil, and each nest contains 3-4 brood cells. Once a brood cell is excavated, the inside is moistened and compacted by the female using her pygidial plate. It is then lined with a thick, white, waxy secretion that has a cheesy aroma. This lining has been studied in the closely related *Anthophora abrupta;* it is produced in the female’s Dufour’s gland (which occupies nearly 50% of her abdomen) and consists mostly of triglycerides which are mixed with salivary secretions and converted to solid diglycerides during cell construction (31). The specific production of this brood cell lining is thought to be highly specialized for larval consumption, as it is energy rich and converted upon secretion in the brood cell to a more digestible form (31, 32). The lining is eaten by the larva as food just after the pollen provision is consumed, and just before diapause as a prepupa (Fig 1A).

**Figure 1-.**
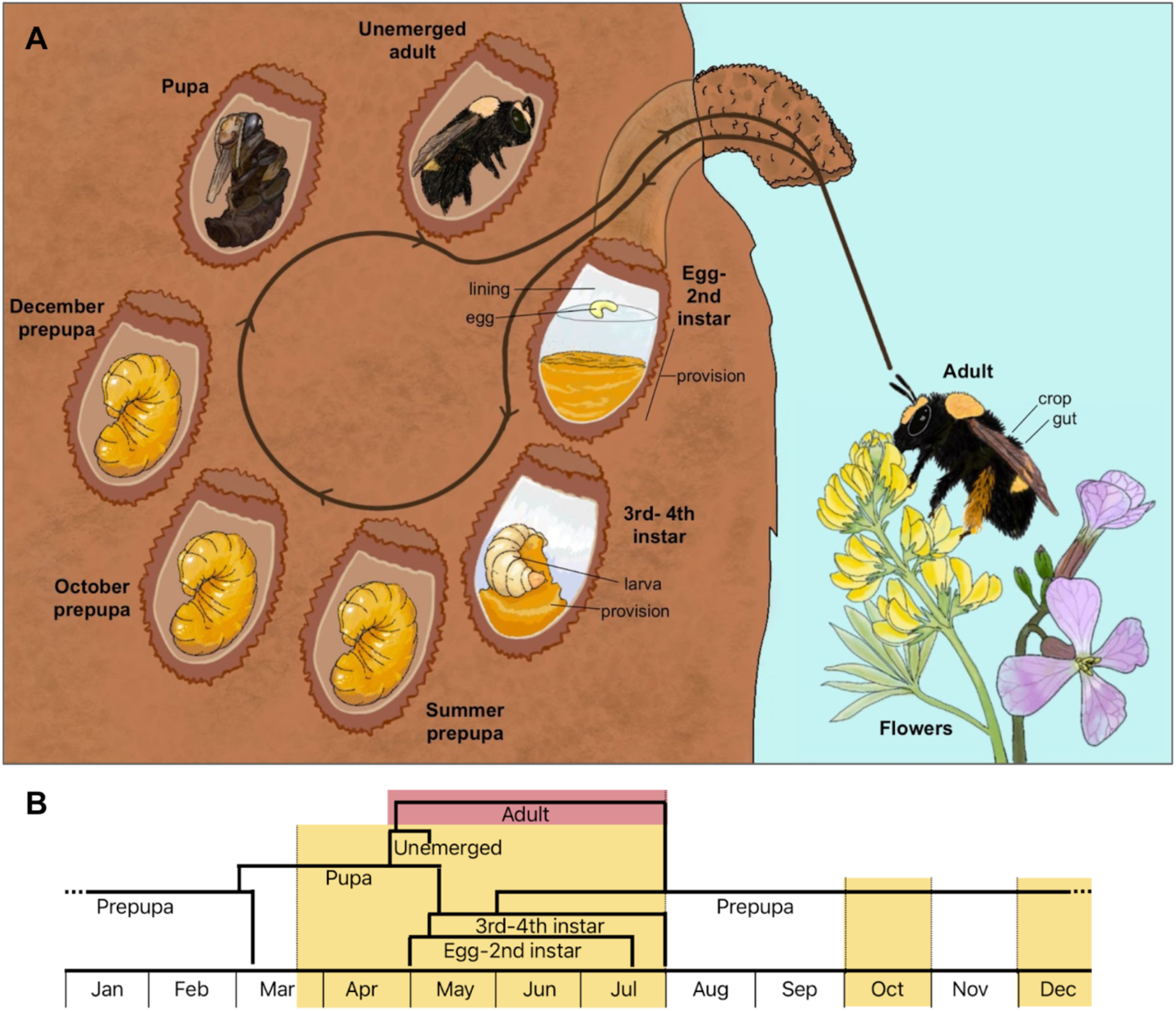
Life cycle of *Anthophora bomboides* indicating sampled stages. **(A)** Full circle represents one year, with labeled brood cells illustrating stages that were sampled. The egg-2nd instar stage includes pollen provisions that contained liquid nectar, only provision was sampled. The 3rd-4th instar stage contained pollen but no nectar and was separated into larva and provision samples, summer prepupae (collected pre-Aug) had recently eaten the cell lining, defecated, and turned yellow-orange. Overwintering prepupae collected in October and December are categorized as such. Pupal and unemerged adult stages were collected in spring; the latter were distinguished by complete development of hairs. Active foraging adults were collected and dissected for crop and gut samples. Stages are listed stepwise as they occur for one bee. Black text and lines indicate which parts of the stage were used for further analysis, excepting “lining” and “egg” which are labeled for illustrative purposes. Absence of lines indicate that the whole bee was used as the sample for that stage. **(B)** At the population level, some stages overlap in time. Highlighted sections indicate months when sampling occurred. Yellow indicated that sampled stages occur within the brood cell, red indicates that sampled stage occurs outside of the brood cell. Illustration by S. Christensen.

*Anthophora bomboides* actively nests from late April to July and is polylectic, but at our sampling locations, prefers to nectar on radish (*Raphanus sativus*) while collecting pollen from lupine (*Lupinus arboreus*) - which are present for the entirety of the nesting season (Fig. 1). Bees also foraged on *Eschscholzia californica*, *Cakile maritima*, and *Carpobrotus edulis*. Building and provisioning a single brood cell takes 2-3 days (27). Pollen is provisioned first and placed in the bottom of the cell (∼5mm thick) then is covered with a layer of nectar (∼630ul) making the overall brood provision initially very liquid (Fig. 1) (27).

After hatching from the egg, young bee larvae (1st-2nd instars) consume first the nectar portion of the provision, followed by the pollen (3rd-4th instars), and finally the cell lining, upon which the white larva turns to a yellow prepupa. The provision is consumed within three weeks, the lining in the following two days; this occurs from late April to July, and in the population, these stages are overlapping. The larva then defecates and diapauses as a prepupa from fall through early spring, and do not spin cocoons. In early spring (Mar-Apr) the prepupae conclude diapause and pupate, and male and female adults emerge between late April and mid-May (Fig. 1) (27).

#### Study Sites

Two nest sites on the California coast were sampled in this study: McClure’s Beach (Point Reyes National Seashore, Marin County, CA, USA) and Bodega Head (Bodega Bay Marine Lab, Sonoma County, CA, USA). The sites are 9.8 miles apart (as the crow flies) and separated by a 5-mile open stretch of ocean (Bodega Bay) between Tomales Point and Bodega Head. McClure’s Beach comprises roughly 2000-3000 nests estimated in early June of 2021 (estimated by grid count on image)-which would indicate around 1000-1500 or more active females. Both nesting sites have been active for at least 4 decades if not longer (27). At both sites, the nests are closely aggregated (within centimeters) but are not connected underground and are not re-used year to year. Because of the largely Mediterranean climate along the coast, the nesting period is warm and quite dry, but winters are wetter and cooler.

### *Anthophora bomboides* brood cell microbial communities are distinct from the environment and dominated by Actinobacteria and *Moniliella spathulata* yeast

To characterize the composition of the microbial communities and associated environments (Fig. 1, Fig. S1), we used Illumina amplicon sequencing of the V5/V6 region of the 16S rRNA gene and ITS gene for bacteria and fungi, respectively. For both bacteria and fungi, brood cell microbial composition was distinct from environmental sources and from adult gut samples (Fig 2C,D) and dynamic through brood cell development (Fig. 5A,B, Fig S2). Actinobacteria were predominant during all stages of brood cell development in significantly higher relative abundance compared to samples outside of the brood cell (K-W χ²= 37.6, df = 1, p = 8.7e-10; Fig. 2A). All samples contained Actinobacteria, but brood cell samples had 84.9% (n=69, sd=17.9) relative abundance, adult bee samples had 29.9% relative abundance (n=2, sd=12.7), and environmental samples had 18.8% relative abundance (n=15, sd= 20.4).

**Figure 2-.**
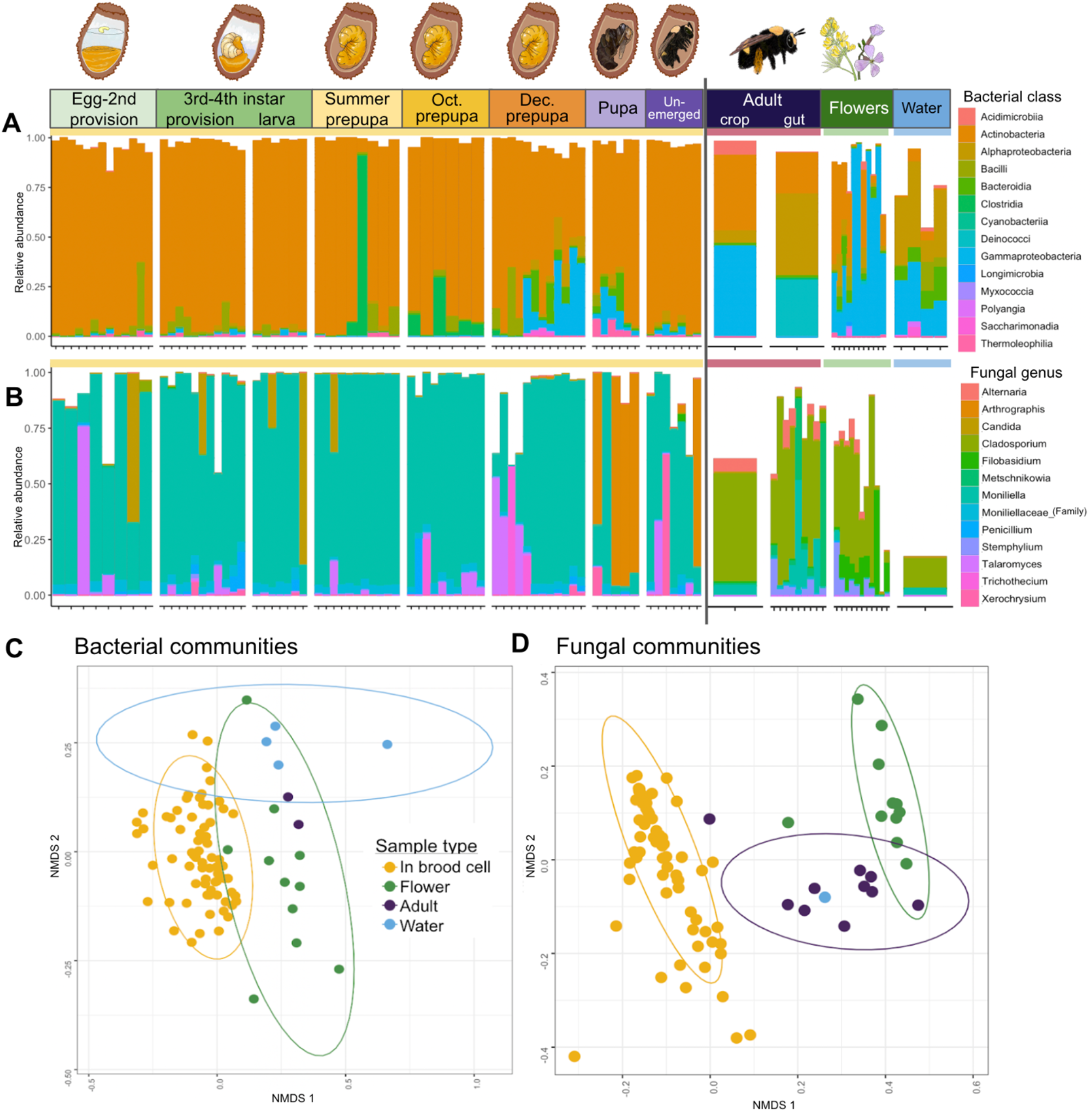
*Anthophora bomboides* brood cell samples are dominated by Actinobacteria and *Moniliella,* differing significantly from adults and environment. **A,B)** Stages arranged in order of development, followed by environmental samples (flowers and water). Dirt samples yielded no sequences after QC and filtering, see Supplemental figure 3. Each vertical bar represents one sample. Yellow horizontal bar indicates that those stages/samples occur within the brood cell, red indicates adults collected outside of the brood cell, green indicates flower samples, and blue indicates water samples where adult bees collect water for nest construction. Black vertical line separates inside (left) versus outside (right) of the brood cell. The same samples were sequenced for both bacteria and fungi, but discrepancies occur when a sample was filtered out due to low read count in one sequencing run but not the other, such as the adult gut sequencing well for fungi and poorly for bacteria. **(A)** Top 500 bacterial ASVs, colored by class. Actinobacteria (orange) dominate within the brood cell. Kruskal-Wallis χ²= 37.6, df = 1, p = 8.7e-10 comparing relative abundance of Actinobacteria from inside brood cell samples (mean 84.9%; yellow bar; left of line) to outside brood cell samples (mean 20.1%; red, green, blue bars; right of line). N=86 samples. **(B)** Top 15 fungal ASVs, colored by genus. *Moniliella* (light blue) dominate within the brood cell. Kruskal-Wallis χ² = 43.9, df = 1, p = 3.3e-11 comparing relative abundance of *Moniliella spathulata* from inside brood cell samples (mean 72.3%; yellow bar; left of line) to outside brood cell samples (mean 5.4%; red, green, blue bars; right of line). N=93 samples. **C,D)** For both plots, color indicates sample type. Yellow corresponds to samples from inside the brood cell, green corresponds to flower samples, purple to adults that were free-flying outside of the brood cell, and blue to water samples. **(C)** NMDS of weighted Bray-Curtis (BC) distance for bacterial communities (stress = 0.19). PERMANOVA with sample type as a predictor R^2^= 0.074, F=2.2, p<0.001. **(D)** NMDS of weighted Bray-Curtis distance for fungal communities (stress=0.1). PERMANOVA with sample type as a predictor R^2^= 0.25, F=10.12, p<0.001.

To determine if Actinobacteria are specifically affiliated with *A. bomboides* we compared relative abundance of Actinobacteria ASVs (489 total) in brood cell and environmental samples (Fig. S4). With a 0.1% detection threshold, 60% were found exclusively in the brood cell samples (294 ASVs), 17% were found in both brood cell and environmental samples (64 ASVs) and 25% were only detected in environmental samples (123 ASVs). Of the 64 ASVs shared between brood and environmental samples, 54% were found in only one environmental sample, mostly flower samples likely visited by foraging *A. bomboides* (*Raphanus sativus, Erigeron glaucus*). These 64 shared ASVs make up on average 9% of the reads in environmental samples (sd= 13.56). Six Actinobacteria ASVs are shared between the adult bee gut samples and the brood cells, comprising an average of only 1.6% of the reads within the brood cell samples (sd= 1.85).

Fungal communities within brood cells were dominated by *Moniliella spathulata* across all stages except the pupal stage, where *Arthrographis* is dominant (Fig. 2). *Moniliella spathulata* was detected in every brood cell where it comprised 72.3% of sequences (n=71, sd=29.6), significantly greater than in adult bees or environmental sources (Fig 2B, K-W χ² = 43.9, df = 1, p = 3.3e-11). While *M. spathulata* was detected in all but one adult GI tract sample, its average relative abundance in this habitat was 10.9% (n=10, sd=16.1). Three environmental samples (*Raphanus sativus* bulked 15x flowers, *Carpobrotus edulis* flower, water sample) contained *M. spathulata;* the mean relative abundance in environmental samples was 0.7% (n=12, sd=1.57).

### *Anthophora bomboides* brood cells host a core microbiome composed of select Actinomycete genera and *Moniliella spathulata* yeast

We next sought to evaluate the presence and composition of a core microbiome in *A. bomboides* brood cells using the 16S rRNA and ITS gene amplicon data. Bacterial core was defined at the genus level for samples inside the brood cell, because the genera seemed to remain quite consistent despite diversity at the ASV level. At a prevalence of 65% and detection threshold of 0.1%, six genera (*Streptomyces, Arthrobacter, Nocardioides, Mycobacterium, Pseudarthrobacter,* and *Rhodococcus*) comprise a bacterial core. At the stricter prevalence cutoff of 90%, *Streptomyces* and *Nocardioides* remain core genera (Fig. 3A). Regardless of the specific numerical cutoff, the top 8 genera that could constitute the core are all Actinobacteria (marked with * on Fig. 3A).

**Figure 3-.**
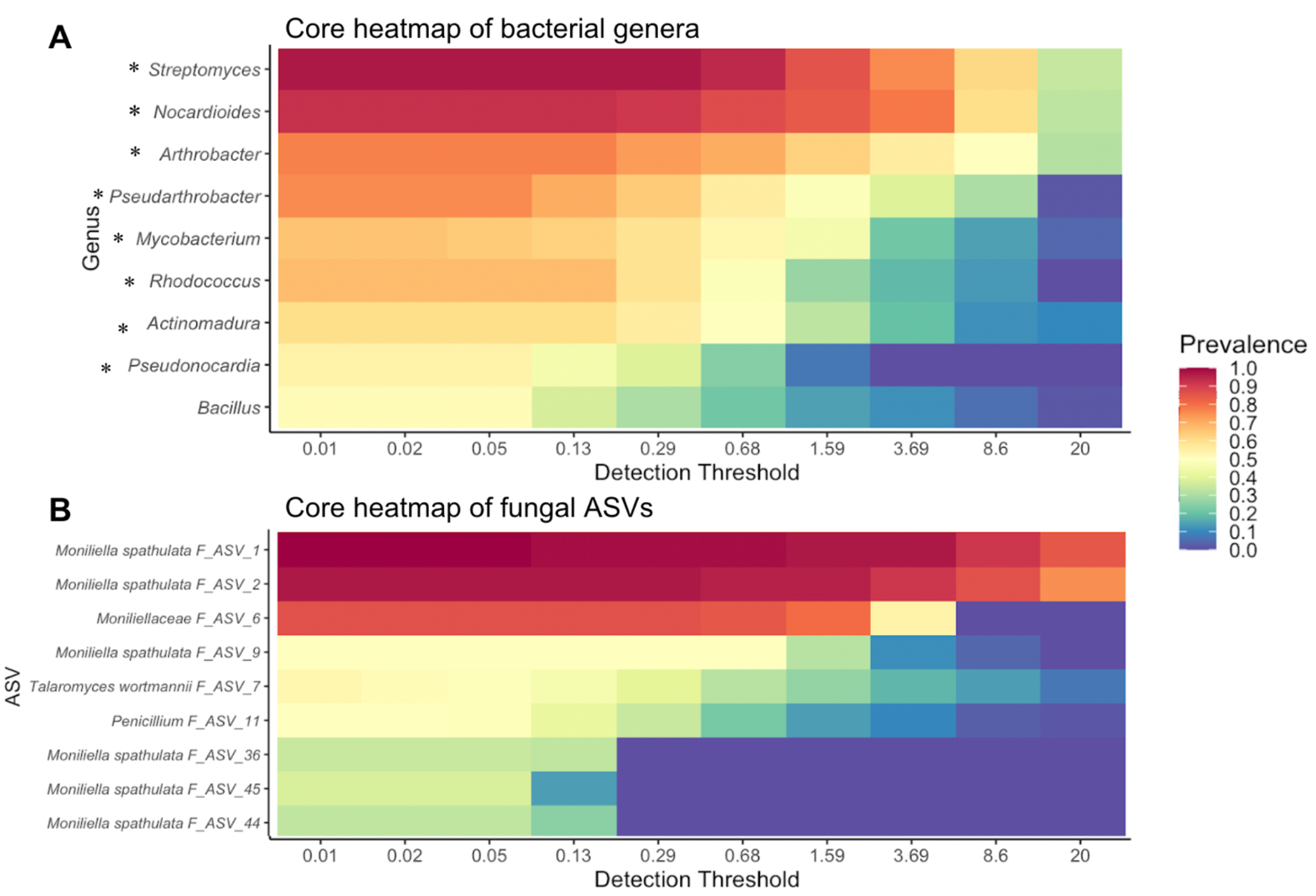
*Anthophora bomboides* brood cells host a core microbiome composed of Actinomycetes and *Moniliella*. Core microbiome heat maps for bacterial genera (A) and fungal ASVs (B), indicating prevalence at increasing detection thresholds. Prevalence is the proportion of samples containing the indicated taxa, detection threshold is the minimum relative abundance that needs to be present in a sample for it to be counted. Together, these separate core taxa (high prevalence and abundance) from other taxa. Top taxa arranged in decreasing order down the y axis, prevalence for each taxa is indicated by color, with 1 (dark red) meaning that the taxa is present in all samples, and 0 (dark blue) indicating it is present in none of the samples at each detection threshold (x axis, % of reads). **(A)** All bacterial genera with an asterisk (*) belong to Actinobacteria class. **(B)** Fungal ASVs labeled at the most specific assigned taxonomic level, see Fig. S6 for relative abundance of fungal ASVs 1&2.

The fungal core microbiome was defined at the ASV level for samples inside the brood cell. Two ASVs in particular, ASV_1 and ASV_2, both assigned to *Moniliella spathulata* are present in 97% of samples at a detection threshold of 0.5%, and still 88% of samples at a detection threshold of 5% (Fig. 3B). ASV_6, also belonging to Moniliellaceae, could be considered core at lower thresholds, but no other ASVs approach inclusion in the fungal core within the brood cell (Fig. 3B).

**Figure 4-.**
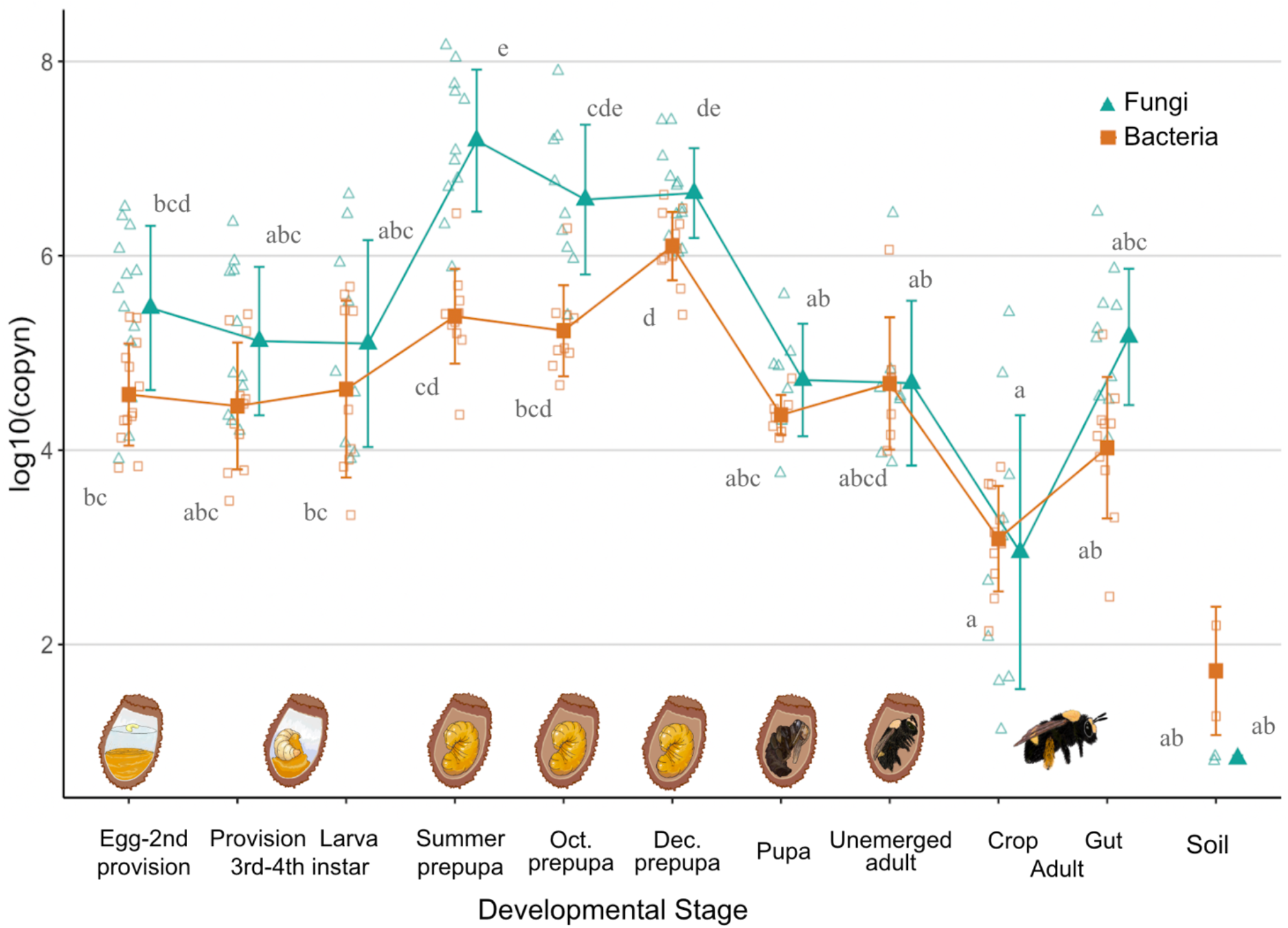
*Anthophora bomboides* microbial copy number is highest in diapausing prepupae. Filled points represent the mean of the log10(copy number) by developmental stage, error bars +/-1 SD log(copy number) of each developmental stage for both bacteria (orange squares) and fungi (teal triangles). Smaller open points represent individual samples. Bacterial copy number has been adjusted to remove non-bacterial reads, as determined by amplicon sequencing with identical primers. Kruskal-Wallis test indicates significant differences in bacterial and fungal copy number based on bee developmental stage, (Bacteria-KW χ^2^= 66.7, df = 10, p-value = 1.9e-10; Fungi-KW χ^2^ = 67.0, df = 10, p-value = 1.6e-10). Lettering indicates differences via Dunn’s multiple comparisons; above for fungi and below for bacteria (p<0.01).

### Microbial abundance peaks in during bee diapause

To quantify abundance of bacteria and fungi throughout the bee life cycle, we conducted qPCR using the same bee samples as above. Bacterial copy number increased through bee development (Fig. 3; KW χ^2^= 66.7, df = 10, p= 1.9e-10). Specifically bacterial copy number increases through larval development, peaking during December, mid-diapause, and decreasing after pupation; bacterial copy number in December prepupae is 33 times higher than in egg-2^nd^ instar (1.3e6, 3.7e4).

Fungal qPCR was conducted using FungiQuant (33). Fungal copy number also changed through development (Fig. 3, KW χ^2^ = 67.0, df = 10, p = 1.6e-10) increasing 52x between egg-2^nd^ instar stage and Summer prepupae (2.8e5, 1.5e7), coinciding with consumption of the brood cell lining and defecation. Fungal copy number remained high through December before dropping by a factor of 83x between December prepupal stage and pupal stage (4.3e6, 5.2e4).

### Stages within the brood cell have shifting bacterial and fungal communities, *Streptomyces* dominates in overwintering stages

To determine whether the communities shift at finer taxonomic scales within the brood cell during development, we used the 16S rRNA and ITS amplicon sequencing data to compare bacterial and fungal communities at different stages inside the brood cell. Microbial communities within the brood cell consistently contain a high proportion of Actinobacteria and *Moniliella spathulata*, but the relative composition changes between stages after the Summer prepupa stage (p<0.05, Fig. 5A). Fungal community composition was overall consistent between stages, but was distinct in Summer prepupae and pupae (p<0.05, Fig. 5B).

**Figure 5-.**
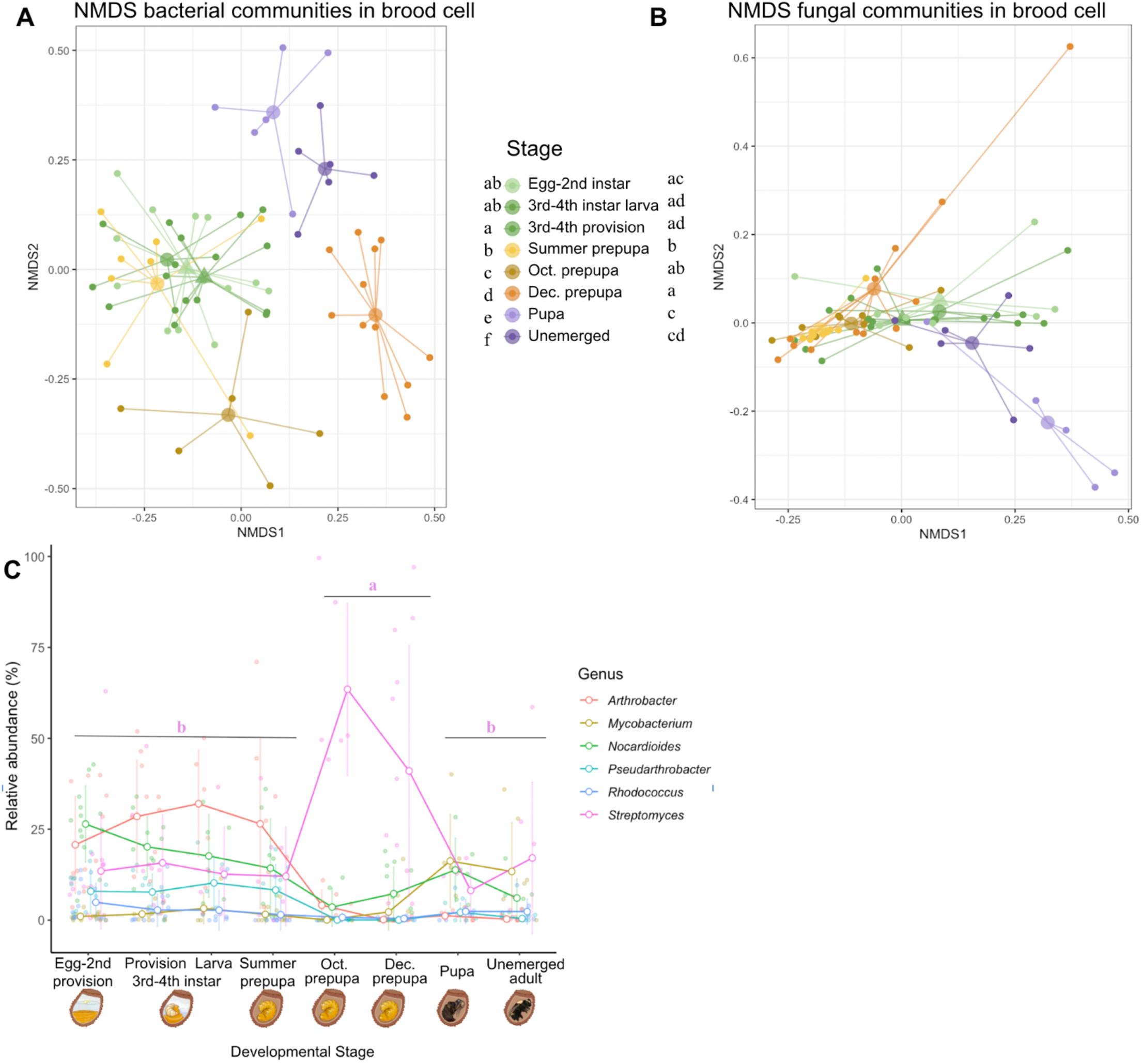
Bacterial and fungal communities shift with bee development, with increased *Streptomyces* abundance in overwintering stages. **(A)** NMDS of Bray-Curtis distance with color indicating stage of brood cell development. Larger semi-transparent dots indicate centroids, with lines from centroid to each point in the group. Triangular centroids indicate provision samples. Bacterial-global PERMANOVA shows significant difference between stages (R^2^=0.25, F=2.87, p<0.001). Pairwise PERMANOVA of stages (FDR<0.05) indicated with lettering on left side of figure key. **(B)** Fungal-global PERMANOVA shows significant difference between stages (R^2^=0.29, F=3.62, p<0.001). Pairwise PERMANOVA of stages (FDR<0.05) indicated with lettering on figure key. **(C)** Relative abundance (% of total) of top 6 bacterial genera across bee developmental stages. Six genera comprise the brood cell core at min prevalence of 65% of samples. Larger open circles represent mean relative abundance of the genus at the indicated developmental stage and are connected by lines. Error bars are (+/-1 SD). Each smaller shaded point represents the relative abundance of the corresponding genus in one sample. *Streptomyces* relative abundance varies through development (Kruskal-Wallis χ *^2^*= 15.9, df = 2, p = 0.0003) and is greatest during overwintering (Oct.-Dec.) as compared to summer (Egg-Summer prepupa; Dunn’s test adj. p<0.001), or spring stages (Pupa-Unemerged adult; Dunn’s test adj. p<0.01).

We further examined how the relative abundances of the top six bacterial core genera change with developmental stage. The most abundant genus, *Streptomyces*, peaks in abundance in overwintering prepupae, with average relative abundance of 48.5% (sd =35.2, during October and December). *Streptomyces* relative abundance in the overwintering stages is significantly greater than summer (egg through Summer prepupae, avg. 13.7% sd=13.8, p = 6e-4) or spring (pupae and unemerged adults, avg. 13%, sd= 16.1, p=0.003) stages by Dunn’s test (Fig. 5C).

### *Streptomyces* isolated from brood cells inhibits growth of filamentous fungi

Fungal pathogens can thrive in the wet conditions of the overwintering period, which also coincided with peak abundance of *Streptomyces.* To test the hypothesis that *Streptomyces* can inhibit the growth of filamentous fungi, we used a plate-based co-inoculation assay (see Methods). We examined whether *Streptomyces* isolates from *A. bomboides* brood cells (BH34, BH55, BH97, BH104) could inhibit the growth of *Ascosphaera apis*, a devastating pathogen of bee brood, as well as *Thelonectria*, which we isolated from an infected pupa of *A. bomboides. Streptomyces* isolates from the brood cells were able to inhibit both fungi. Although *Streptomyces* strains varied in their fungal growth suppression, *A. apis* was significantly inhibited by both BH34 (p<0.05) and BH97 (p<0.05) and *Thelonectria* was significantly inhibited by BH34 (p<0.01) and BH55 (p<0.05) on day 7 of co-inoculation (Fig. 6 A,B,E). As BH34 was able to inhibit both pathogenic fungi, we then tested whether it would also inhibit *Aspergillus flavus*-the generalist bee pathogen, or *Moniliella spathulata*-the core fungal taxa, using the same methods. We found that BH34 was also able to significantly inhibit the growth of both *Aspergillus flavus* (p<0.01, day 4) and brood cell isolated *Moniliella spathulata* (p<0.01, day 7; Fig. 6C, D).

**Figure 6-.**
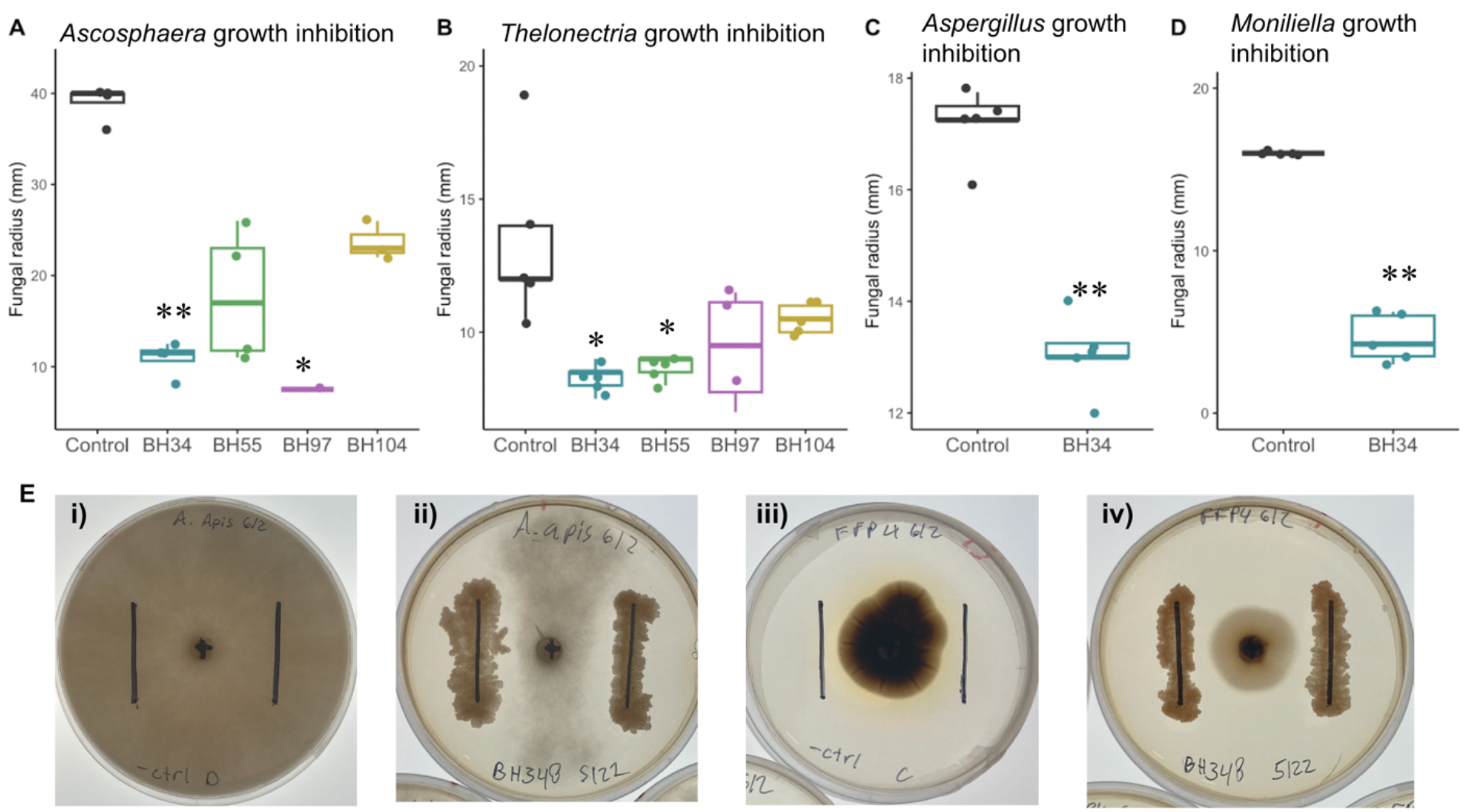
*Streptomyces* isolated from *A. bomboides* brood cells inhibit growth of filamentous fungi. Inhibition is shown by decrease in radius of fungi (mm, y axis) on plates when co-inoculated with isolates of *Streptomyces* (BH104, BH34, BH55, BH97) (x axis, colors) from brood cells as compared to control (black). *Streptomyces* isolates plated 10 days prior to inoculation with 6mm diameter fungal plug. Comparisons to control by Dunn test, p-val adjust with Bonferroni (p<0.05 = *, p<0.01= **). (**A-D**) Center lines correspond to the median, boxes define interquartile range (IQR), and whiskers extend +1.5 IQR. Note difference in scale of y axes. **(A)** Inhibition of *Ascosphaera apis,* a common pathogen of bee brood, on day 7 of co-inoculation. Data represents 16 plates. BH34 and BH97 significantly reduced the radius of *A. apis.* Note 40mm radius was the edge of the plate. Kruskal-Wallis *X^2^* = 12.3, df = 4, p-value = 0.014. **(B)** Inhibition of *Thelonectria,* a potential pathogen isolated from an infected pupal cell, on day 7 of co-inoculation. Data represents 24 plates. BH34 and BH55 significantly reduced the radius of *Thelonectria*. Kruskal-Wallis *X^2^*= 14.5, df = 4, p-value = 0.005. **(C)** Inhibition of *Aspergillus flavus*, generalist bee pathogen, by BH34 on day 4 of co-inoculation. Data represents 10 plates. Kruskal-Wallis *X^2^*=6.9, df=1, p-value= 0.009. **(D)** Inhibition of *Moniliella spathulata*, ubiquitous brood cell yeast, by BH34 on day 7 of co-inoculation. Data represents 10 plates. BH34 significantly reduced the radius of *Moniliella spathulata.* Kruskal-Wallis *X^2^*= 7.3, df = 1, p-value = 0.007. **(E)** Plate images of fungi on day 7 of growth, when grown alone and with *Streptomyces* isolate BH034. i) *A. apis* negative control, ii) *A. apis* with *Streptomyces* isolate BH034, iii) *Thelonectria* negative control, iv) *Thelonectria* with *Streptomyces* isolate BH034. Notable is not only the reduction in radial growth but also qualitative density and color of fungal hyphae. Images were taken on backlit LED screen to ensure identical lighting.

### Sugar alcohol profiles distinguish developmental stages and coincide with changes in fungal abundance

The genus *Moniliella* is known for its industrial production of sugar alcohols and energy storing carbohydrates such as erythritol, glycerol and trehalose (34, 35). In insects, trehalose is protective against environmental stress, such as temperature extremes, dehydration, oxidation and starvation (36, 37). To determine if *A. bomboides* stages exhibit changes in sugar and sugar alcohol composition in development that may coincide with proliferation of fungi, of which the vast majority are *Moniliella*, we analyzed 4^th^ instar larvae (before fungal proliferation), prepupae (highest fungal abundance) and pupae (when fungal abundance drops, and *Moniliella* is replaced by *Arthrographis*; see Figs. 2, 4) to determine their composition of sugar and sugar alcohols (SSA) via HPLC-CAD. We found that the stages with high fungal abundance (prepupal, diapausing stages) have distinct SSA composition as compared to 4^th^ instar and pupal stages (p<0.01 by pairwise PERMANOVA; p-values FDR corrected, Fig. 7A). Specifically, glucose/sorbitol and fructose decline in relative abundance as bees develop from late-stage larvae to prepupae, while the disaccharide trehalose increases to high relative abundance in Summer prepupae, coinciding with peak fungal abundance (*Moniliella*). Trehalose remains the SSA with the highest relative abundance throughout diapause (Fig. 7B).

**Figure 7-.**
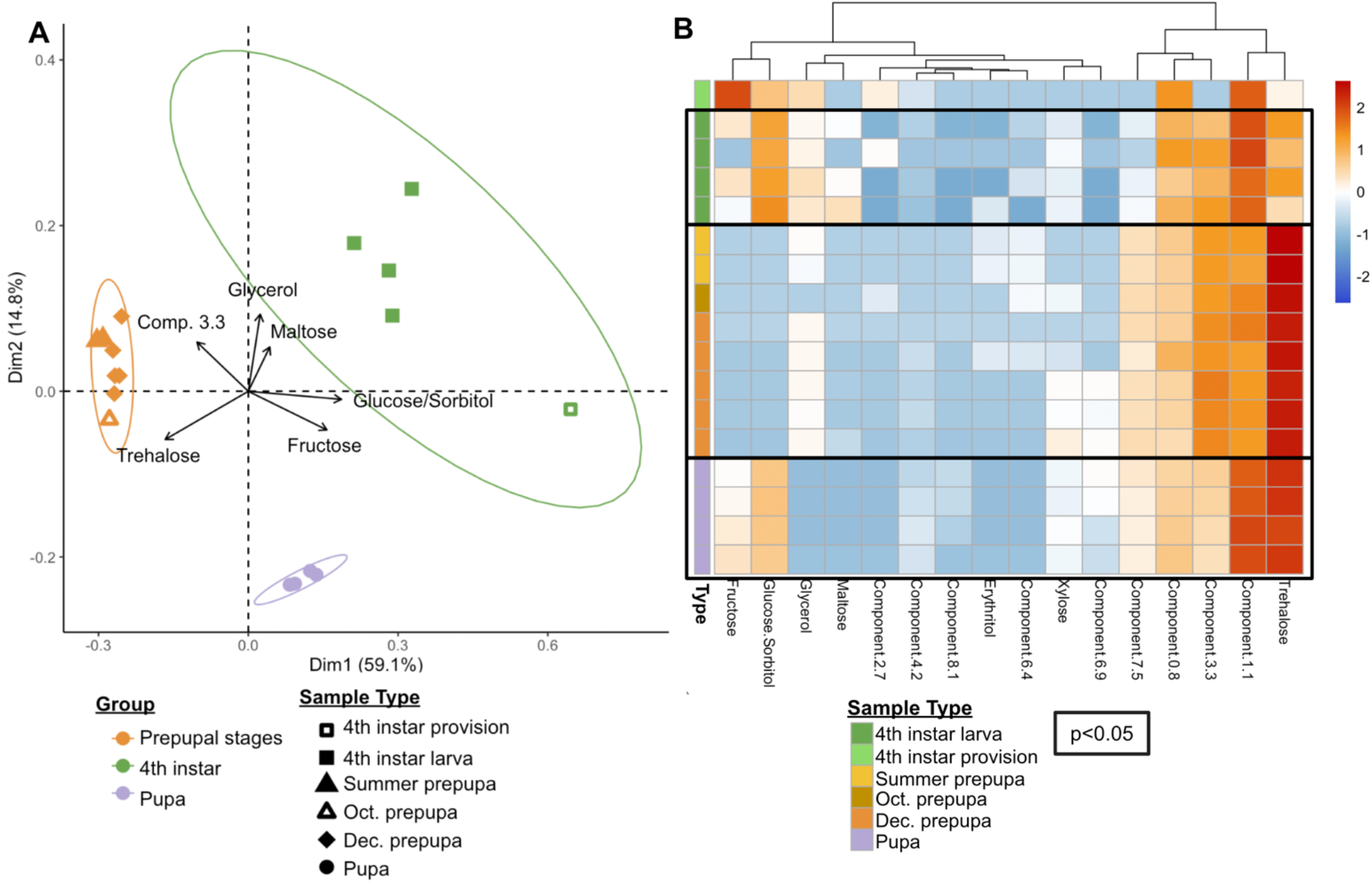
Sugar /sugar alcohol (SSA) composition differentiates bee developmental stages. **(A)** SSA composition of late larvae through pupal stages mapped by PCA, colored by stage. Ellipses indicate 95% confidence for sample groups. Axes labeled with % variance explained. Biplot of the 6 most influential SSA components; longer arrows indicate greater influence on sample separation, direction indicates the alignment with the mapped PCs. Glucose and sorbitol had overlapping and thus indistinguishable peaks. **(B)** Composition of individual SSAs (columns) of each sample (rows). Samples grouped and colored by stage in the first column. SSA data was Hellinger transformed and scaled to sample (relative abundances) with red indicating high relative abundance of that component (column) for that sample (row) and blue indicating low relative abundance within the sample. Black boxes indicate significant clustering of sample compositions (p <0.05) via hierarchical clustering (multiscale bootstrap resampling).

## Discussion

### Unique core

Our study provides a detailed characterization and demonstration of the potential for complex interplay of insect development with microbiome composition and abundance. In contrast to previously studied solitary bee species, we identified a complex core microbiome in the brood cells of the solitary bee *Anthophora bomboides* which increases in abundance with larval diapause and persists across life stages. Our findings in *A. bomboides* contrast with previously characterized solitary bee microbiomes in a few key ways. First, most previous work on solitary bees suggests that their microbiomes tend to be acquired from, and thus mirror, their immediate environment (16, 19, 38–40), whereas *A. bomboides* microbiomes are distinct from the environment. Whereas fungal associates are often left out of most bee microbiome characterizations, (41) or found to decrease in diversity and abundance over time (42), *A. bomboides* hosted abundant and distinct fungi in brood cells. Even at the earliest stage, provisions had fungal composition that was dominated by *Moniliella spathulata* (Fig. 2). The only other documented persistent mutualistic interaction between bees and fungi has been in social stingless bees, where yeasts are persistently found in the colony and produce a steroid required for larval development (43). In *A. bomboides*, both bacterial and fungal brood cell microbiomes were distinct from environmental sources including floral resources and water used to soften soil for brood cell construction. Minimal bacterial ASV sharing between brood cells and flowers further suggests that the mode of *A. bomboides* microbiome acquisition is distinct from what has been characterized in other solitary bees.

Secondly, the microbiome described here is complex and nearly all brood cells are dominated by 6 core bacterial genera and a single fungus. In previous work on solitary bees, brood cell bacterial microbiomes are highly variable among individuals or composed of more limited taxa (18, 19, 40, 44–46). Most commonly in brood cells with consistent microbial communities, lactic acid bacteria are dominant in the provisions, but disappear before pupation (18, 19). The loss or extreme reduction of bacteria upon pupation and emergence as adults, combined with evidence suggesting strain sharing between flowers and hosts points to annual environmental acquisition (18, 38, 47). This acquisition can lead to divergence in microbiomes depending on habitat, as seen in *Megachile* gut bacteria as it invades new areas and shares microbes with co-foraging bee species (48). In comparing the bacteria associated with the provisions and larvae of four solitary species, it was found that there are species-specific patterns and even prevalent members, but high diversity and fluctuation, attributed to the lack of vertical transmission and solitary lifestyle (46). Another recent study describing bacteria associated with the provisions and larvae of four solitary species found that bee species host distinct communities but also documented high species diversity and turnover among nests within a species (46).

Taken together, the lack of evidence for environmental acquisition and the consistency of bacterial genera and fungal ASVs in brood cells among sites and years suggests vertical transmission of the microbiome in this solitary bee species but this remains to be experimentally confirmed.

Fungus growing ants and stingless bees both have clear routes of social and vertical transmission of their core associates (49–51). Both are social and maintain ongoing nests year-round to allow horizontal transfer between nest-mates, and both are known (or strongly suspected) to establish new nests with clonal propagations of their associated fungi. Solitary bees in general do not have these clear methods of vertical or social transmission, but nonetheless we found that brood cells had highly consistent bacterial and fungal microbiomes. However, these were not consistent with adult GI tracts, which either did not sequence well (Bacteria), or had stronger resemblance to the environmental communities (Fungi). Another solitary hymenopteran, the beewolf (*Philanthus* spp.), transfers its core brood associate to newly provisioned cells using secretions from a specialized gland (8). The data here suggest that some method of assured transmission is occurring for *Moniliella spathulata*, as nearly every cell from both sampled sites contained the same two ASVs, indicating tight and effective control of transmission. Further work will examine potential sources of this complex microbial community and test modes of transmission.

Third, this microbiome composition and its consistency among brood cells is unique among bees (Anthophila). At the strictest cutoffs used here, *Streptomyces* and *Nocardioides* are core bacterial genera of the brood cell, being present in over 90% of samples (min 0.01% abundance). Additionally, nearly every brood cell (97%) contained *Moniliella spathulata* ASVs 1 and 2, most often as the dominant taxa. The bacterial core taxa described here have not been previously described in bees but instead are in some ways comparable to the widely studied tripartite symbiosis of fungus farming ants (52) where workers host *Pseudonocardia* (29, 53) to protect their highly co-evolved cultivated fungus, which is grown in monoculture (50). However, the data presented here suggest that although *Streptomyces* isolated from *A. bomboides* brood cells suppress the growth of pathogenic fungi, they also suppress the growth of *Moniliella spathulata*, the core fungal taxa. This observation suggests that microbial communities may exhibit small-scale niche partitioning between Actinomycetes and yeasts, or that conditions used in the plate assay result in different dynamics compared to the brood cell. Nevertheless, the function and localization of microbial taxa within the brood cell will require additional study.

### Stage specific vulnerabilities- a role for microbes?

The annual life cycle of many insects correspond to and are influenced by seasonal patterns (54). Insects can enter seasonal diapause in order to conserve energy and survive harsh conditions, but may be more vulnerable to predation or pathogens (55, 56). Though diapausing insects retain innate immunity (57), some insect species show reduced or altered immune response during diapause or overwintering (58, 59). The most common cause of brood mortality in *A. bomboides,* and solitary bees in general, is fungal pathogens (60), which thrive in cool, moist conditions (61) experienced during overwintering. We found that bacteria in general and *Streptomyces* specifically attain the highest abundances in overwintering October and December prepupae; with *Streptomyces* absolute abundance increasing by 46-fold between early provisions and December prepupae. This, combined with the demonstrated ability of brood-isolated *Streptomyces* to inhibit fungal pathogens, suggests a defensive mutualism during bee overwintering. Alternatively, lowered bee immune defenses may allow *Streptomyces* to proliferate, or *Streptomyces* may be responding to season independently of the host, so further experiments are required to demonstrate the hypothesized mutualism. However, other bee species tend to exhibit low or undetectable microbial populations following defecation, suggesting distinct biology in *A. bomboides* that supports bacterial and fungal growth.

Overwintering can also lead to more subtle effects on survival via indirect chill injuries, which can lead to a gradual failure to maintain homeostasis(56) and increased oxidative stress (62). Many insects use antifreeze compounds, such as trehalose and glycerol, to reduce ice crystal formation and stabilize proteins (56). Members of *Moniliella,* the ubiquitous yeast found with developing *A. bomboides*, are best known for their prolific production of sugar alcohols and trehalose (63). In exploring the patterns of both sugar alcohol makeup and fungal abundance across developmental stages, we found that shifts in sugar alcohol composition corresponded to shifts in fungal abundance, with high levels of trehalose occurring during overwintering, when fungal abundance was also at its peak. Although insects are also capable of producing sugar alcohols and trehalose (64), and xerophilic yeasts may produce sugar alcohols to enhance their own survival in habitats with low water activity (65), correlation between fungal abundance and trehalose production raises the hypothesis that microbial symbionts could be involved in the production of compounds related to cold tolerance. As *Moniliella spathulata* is also known to be oleaginous and able to accumulate lipids up to >60% of its dry weight, it is also possible that its role may be related to lipid metabolism (66, 67). These, as well as additional adaptive and nonadaptive hypotheses will require further investigation.

### Why Anthophora?

To date, *A. bomboides* is the only solitary bee which is now documented to associate with a complex core microbiome that increases in abundance throughout development. What traits may support this specialized association, and is this consistent core microbiome unique to *Anthophora bomboides*? We hypothesize that the production of a specialized brood cell lining may be involved in maintaining this association. Some species within the genus *Anthophora* are unique in that their Dufour’s gland secretion has evolved from a thin waterproof cell lining to a thick, energy dense food source for developing brood. This lining has been noted as being highly specialized, similar to royal jelly in honey bees or milk in mammals (31). Adult females produce copious amounts of this secretion, using it to both line the cell and mix into the provision itself, giving the brood cells a distinct cheesy aroma during early development(31).

We hypothesize that this unique supplementation may aid in the maintenance of core microbial taxa, perhaps especially for *Moniliella spathulata*, as this yeast species is lipophilic and can degrade a wide range of over 150 hydrocarbons (67). Some Dufour’s secretions also have antibiotic properties, which, if present here, may exhibit selective pressure in shaping the microbiome. Interestingly, *Megachile* species are known to have hypertrophied Dufour’s glands which produce triglycerides, and their brood cell contents are associated with another member of *Moniliella* (*M. megachilensis*, previously *Trichosporonoides*) (68–70). Actinobacteria are frequently isolated from soil and plants, but not often from flowers, with the exception of this study and a study on *Pulmonaria* nectar, in which 50% of the OTUs recovered belonged to Actinobacteria (71). Investigations of *Pulmonaria* pollination showed that *Anthophora plumipes* was an important pollinator, and separate microbial characterization of adults revealed persistent associations with *Streptomyces* (45). *Anthophora urbana* has also been found to host prevalent fungi, which are present in live brood cells (72). These findings indicate that perhaps solitary bee associations with Actinobacteria and *Moniliella* may not be limited to *Anthophora bomboides,* but will require further investigation.

## Conclusions

In conclusion, we provide direct evidence of a consistent and abundant microbial community in developing solitary bees and antifungal activity of the abundant *Streptomyces,* which suggests a protective symbiosis. Solitary bees are vulnerable to multiple sources of mortality during development, especially during overwintering. Increasing microbial titer during vulnerable life history stages, consistent composition across brood cells, and microbial phenotypes with clear links to bee life history suggest (but do not yet demonstrate) a mutualistic symbiosis. Specifically, two ASVs of the yeast *Moniliella spathulata* were consistently found at both sites and all developmental stages, with abundance corresponding to significant shifts in sugar alcohol composition in overwintering, pointing to a role in cold-tolerance. *Streptomyces* was found to be a potential defensive symbiont, inhibiting a variety of brood-pathogenic fungi, and dominating overwintering stages. These results highlight a few underappreciated aspects of insect-microbe symbiosis: 1) a specialized microbiome can be maintained in the absence of sociality, 2) bacteria and fungi may affect bee biology during diapause, and 3) the mycobiome may be important and likely deserves additional study. Although much work remains to examine the ecology of the bee-microbiome, our study reframes the conditions thought to maintain symbiosis and may stimulate novel research in exploring additional roles of the microbiome that promote the evolution of symbiosis.

## Methods

### Sites

McClure’s Beach is the larger nesting site, with roughly 2000-3000 nests estimated in early June of 2021 (estimated by grid count on image)- which would indicate around 1000-1500 or more active females given females are likely to have started a second nest by that time (27). It is located in a WSW facing eroded bluff made of hard granitic sedimentary substrate (70% sand, 12.2% silt, 14.5% clay) and is ∼150’ from the ocean. The Bodega Head nesting aggregation is far smaller, estimated 250-500 nests; 100-250 active females (June 2021) and located on gently sloping sides and tops of eroded ditches. The substrate is finer and darker (76.5% sand, 9% silt, 14.1% clay). The site is also on the Pacific Ocean, though higher than McClure’s, ∼50’ above sea level. The two sites are 9.8 miles apart, as the crow flies, separated by a 5-mile stretch of open water (Bodega Bay). Soil makeup determined by Cornell Soil Health Lab.

### Weather and Climate

Because of the largely Mediterranean climate, the nesting period is warm and quite dry (10.2/ 22.2C average low/high; ∼1.5” rain total from May-Sept), but winter is wetter and cooler (5.0/15.7C average low/high; ∼18.7” rain total from Nov-Feb) and there are on average 7.3 days where it drops to or below freezing (Nov-Feb). Data from Bear Valley Visitor’s center on Point Reyes via PRISM, averages for 2006-2015.

### Sample collection

Samples were collected from 2021-2023 at Point Reyes National Seashore (permit #: PORE-2020-SCI-0022) and Bodega Head (SCSP permit issued 2/24/2020). Adults were collected in June with a net while foraging or as they emerged from their nests in the early morning. To collect brood cell samples, small chunks of the cliff in the nesting area were separated using a soil knife and rock pick. These were then carefully dissected to separate brood cells, which are distinct and can be entirely removed from the surrounding soil matrix. Brood cells were then carefully opened from the top with sterilized tweezers or scoopulas (70% ethanol). Tweezers and/or scoopula were re-sterilized before being used to remove brood cell contents into sterile tubes and between brood cell samples. Egg-2^nd^ instar stage brood cells have a high proportion of nectar in provisions; in some cases, a pipette was used to transfer these provisions. Upon collection, samples were rated for how ‘clean’ the extraction of provision was (eg. some had more dirt fall in, or a larva was punctured by the tweezers and soil then stuck to it, etc). The developmental stage, location, and date of collection were also recorded. Developmental stages are described in Table 1. Tubes with collected samples were placed immediately into a cooler for transport back to the lab. Some were vortexed in a buffer (Phosphate Buffered Saline, PBS) and plated for bacterial and fungal isolation (Tryptic Soy Agar, TSA and Yeast Media, YM) or plated directly. The samples that were used for DNA extraction were moved directly from the cooler into a −80C freezer until sample processing.

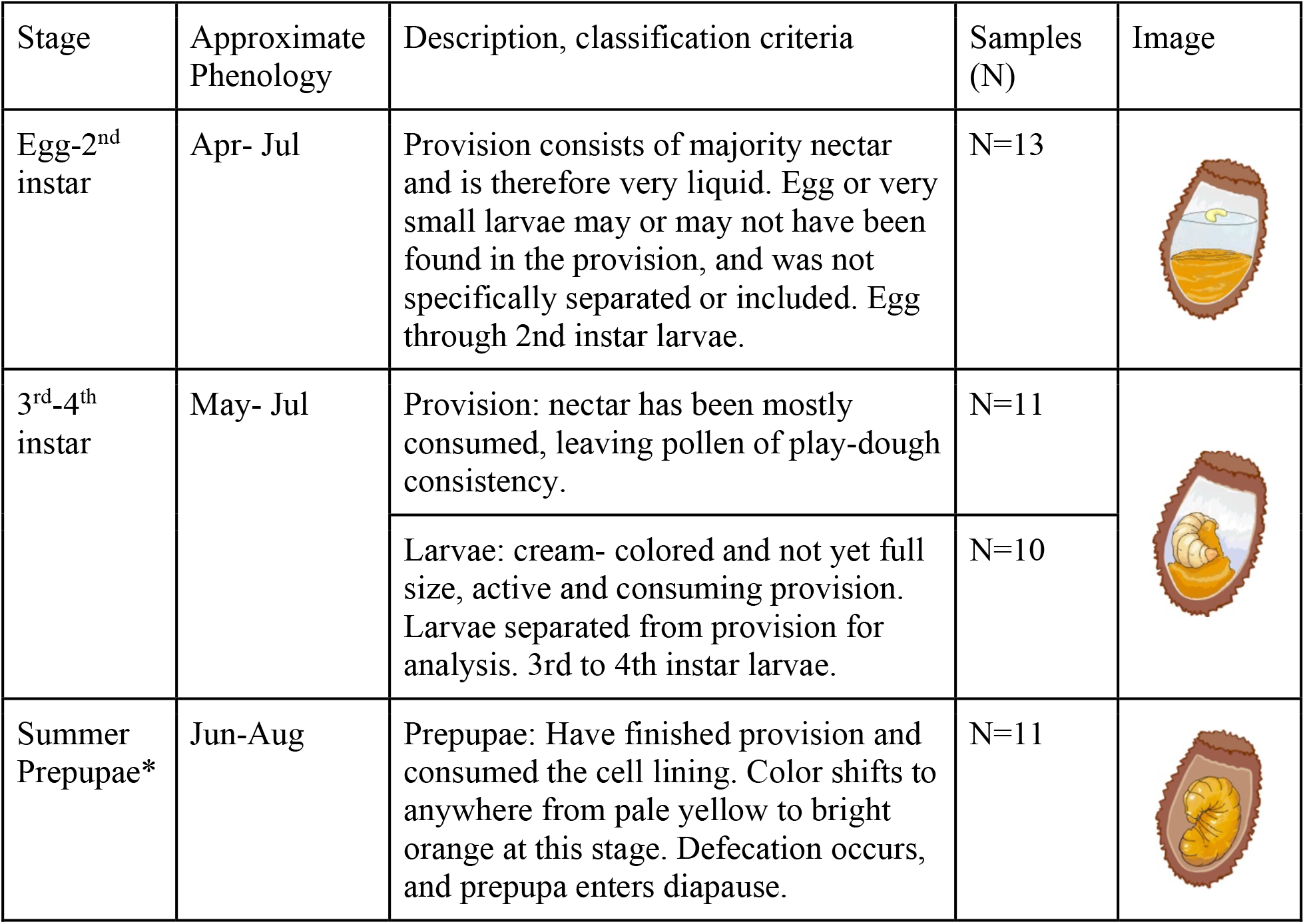
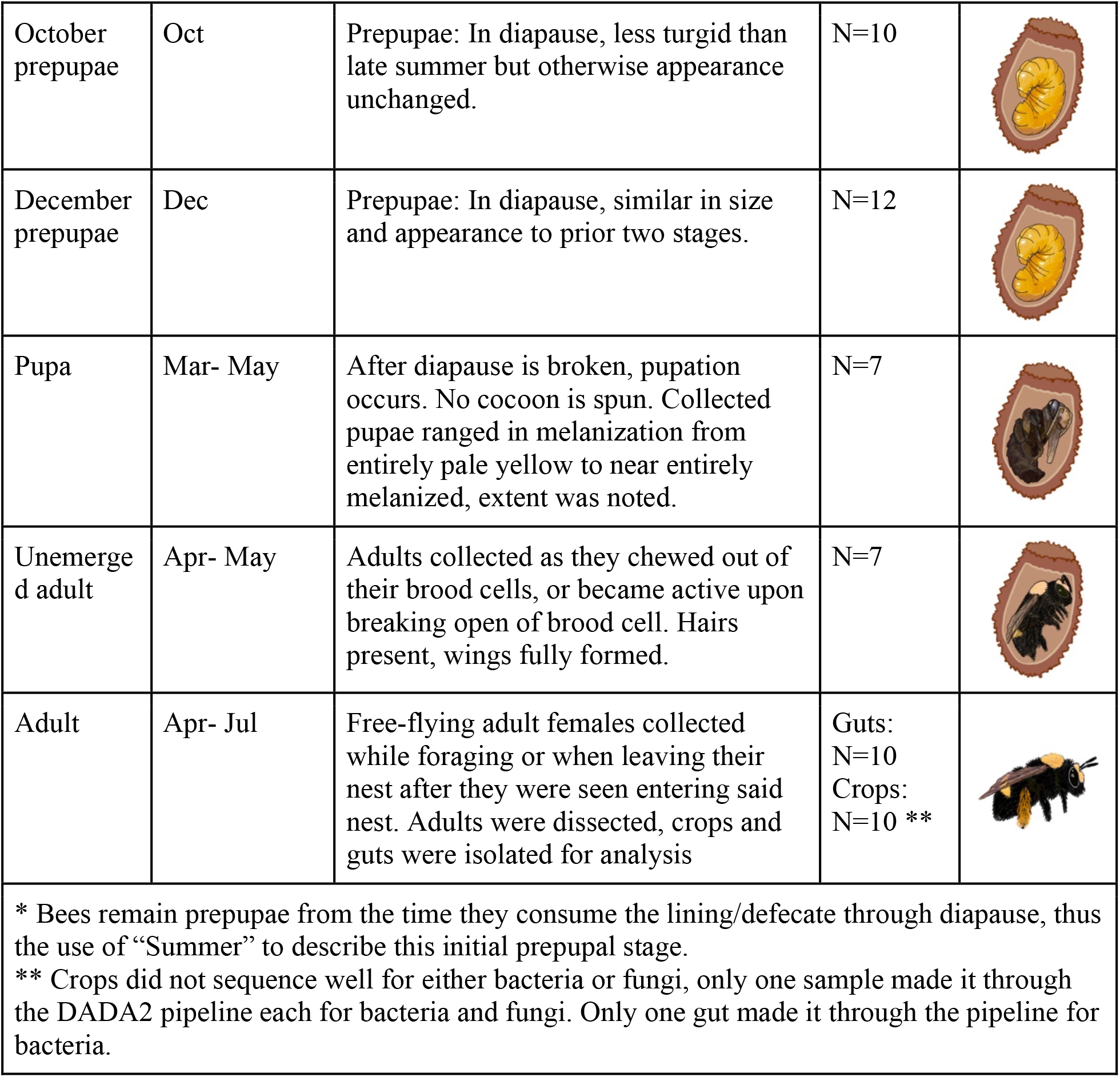
Table 1.

**Table 1**
| Stage | Approximate Phenology | Description, classification criteria | Samples (N) | Image |
| --- | --- | --- | --- | --- |
| Egg-2<sup>nd</sup> instar | Apr- Jul | Provision consists of majority nectar and is therefore very liquid. Egg or very small larvae may or may not have been found in the provision, and was not specifically separated or included. Egg through 2nd instar larvae. | N=13 |  |
| 3<sup>rd</sup>-4<sup>th</sup> instar | May- Jul | Provision: nectar has been mostly consumed, leaving pollen of play-dough consistency. | N=11 |  |
|  |  | Larvae: cream- colored and not yet full size, active and consuming provision. Larvae separated from provision for analysis. 3rd to 4th instar larvae. | N=10 |  |
| Summer Prepupae* | Jun-Aug | Prepupae: Have finished provision and consumed the cell lining. Color shifts to anywhere from pale yellow to bright orange at this stage. Defecation occurs, and prepupa enters diapause. | N=11 |  |
| October prepupae | Oct | Prepupae: In diapause, less turgid than late summer but otherwise appearance unchanged. | N=10 |  |
| December prepupae | Dec | Prepupae: In diapause, similar in size and appearance to prior two stages. | N=12 |  |
| Pupa | Mar- May | After diapause is broken, pupation occurs. No cocoon is spun. Collected pupae ranged in melanization from entirely pale yellow to near entirely melanized, extent was noted. | N=7 |  |
| Unemerged adult | Apr- May | Adults collected as they chewed out of their brood cells, or became active upon breaking open of brood cell. Hairs present, wings fully formed. | N=7 |  |
| Adult | Apr- Jul | Free-flying adult females collected while foraging or when leaving their nest after they were seen entering said nest. Adults were dissected, crops and guts were isolated for analysis | Guts:<br>N=10<br>Crops:<br>N=10 ** |  |
| * Bees remain prepupae from the time they consume the lining/defecate through diapause, thus the use of “Summer” to describe this initial prepupal stage.<br>** Crops did not sequence well for either bacteria or fungi, only one sample made it through the DADA2 pipeline each for bacteria and fungi. Only one gut made it through the pipeline for bacteria. |  |  |  |  |

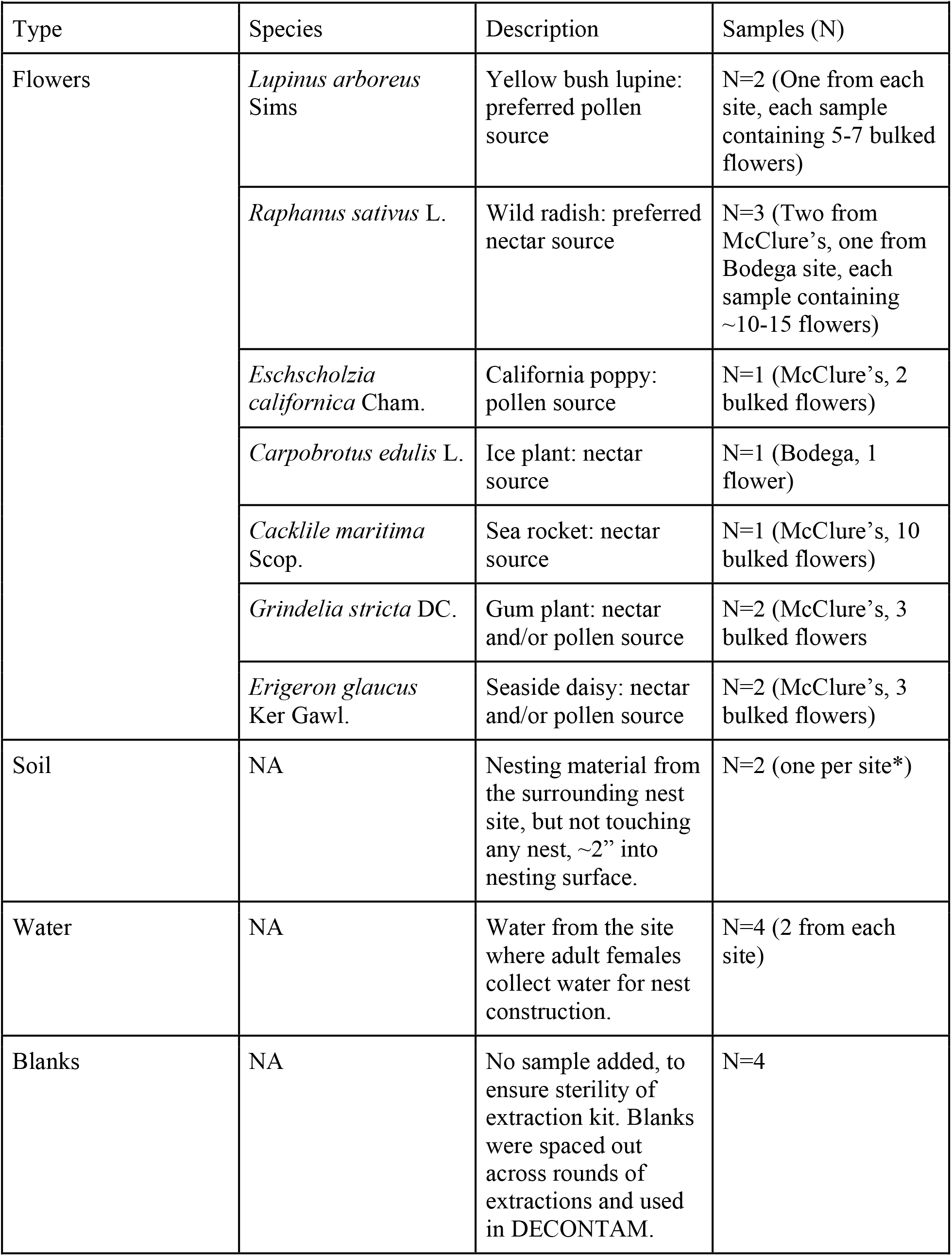
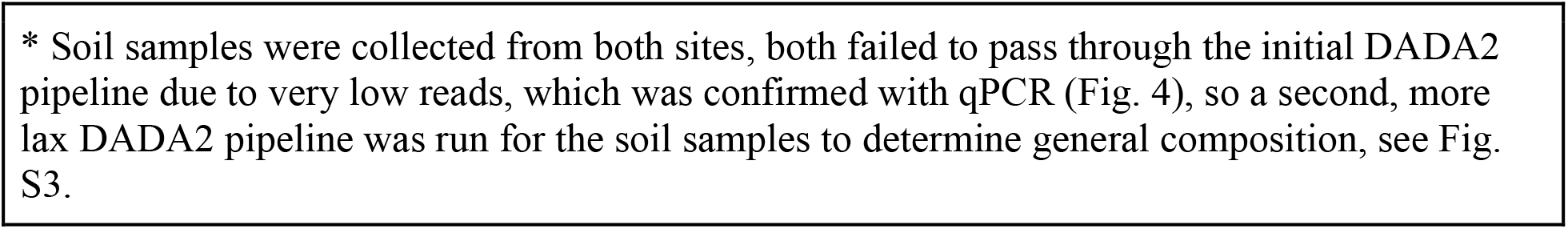
Table 2.

| **Table 2** |  |  |  |
| --- | --- | --- | --- |
| Type | Species | Description | Samples (N) |
| Flowers | *Lupinus arboreus* Sims | Yellow bush lupine: preferred pollen source | N=2 (One from each site, each sample containing 5-7 bulked flowers) |
|  | *Raphanus sativus* L. | Wild radish: preferred nectar source | N=3 (Two from McClure's, one from Bodega site, each sample containing ~10-15 flowers) |
|  | *Eschscholzia californica* Cham. | California poppy: pollen source | N=1 (McClure's, 2 bulked flowers) |
|  | *Carpobrotus edulis* L. | Ice plant: nectar source | N=1 (Bodega, 1 flower) |
|  | *Cacklile maritima* Scop. | Sea rocket: nectar source | N=1 (McClure's, 10 bulked flowers) |
|  | *Grindelia stricta* DC. | Gum plant: nectar and/or pollen source | N=2 (McClure's, 3 bulked flowers) |
|  | *Erigeron glaucus* Ker Gawl. | Seaside daisy: nectar and/or pollen source | N=2 (McClure's, 3 bulked flowers) |
| Soil | NA | Nesting material from the surrounding nest site, but not touching any nest, ~2" into nesting surface. | N=2 (one per site*) |
| Water | NA | Water from the site where adult females collect water for nest construction. | N=4 (2 from each site) |
| Blanks | NA | No sample added, to ensure sterility of extraction kit. Blanks were spaced out across rounds of extractions and used in DECONTAM. | N=4 |

\* Soil samples were collected from both sites, both failed to pass through the initial DADA2 pipeline due to very low reads, which was confirmed with qPCR (Fig. 4), so a second, more lax DADA2 pipeline was run for the soil samples to determine general composition, see Fig. S3.

Flowers, water, and soil were also collected from each location to better understand the environmental microbes that the bees encounter. *Anthophora bomboides* females use fresh water to soften the dirt that they nest in, and often do so in aggregate at a specific location on the spring’s edge. This water was collected into sterile collection tubes from each site at the section that was actively in use by the aggregation at the time of collection. Flowers of each species were collected in June, when adults are foraging, and bulked upon collection into sterile tubes. Nesting substrate, here called “soil”, was collected from within the aggregate nesting area at each site, from within 2” of the cliff surface, several inches from any nests, but within the nesting area. It was collected by chiseling away the exposed surface and then scraping underlying material into a collection tube with a sterile soil knife.

### Sample pre-processing

For collected brood cell samples that had a visible larva and remaining pollen provision (3rd-4^th^ instar), the larvae were separated from the provision and proceeded as two separate samples. Larvae and prepupae were all rinsed gently by pipetting in and removing sterile PBS (2x) before DNA extraction to remove dirt that was introduced in the opening of the brood cells or clinging provision material. For all bee samples, the cleanest samples of each stage, as rated upon collection, were used. Free-flying adult females were dissected, separating the crop and gut as two separate samples per bee. Water samples were centrifuged at 13k for 3 minutes to concentrate the sample for DNA extraction. Top water was removed, and the remaining 100ul was used to resuspend the pelleted solids for extraction. Flower samples were immersed in PBS and vortexed at max speed for 60 seconds to dislodge microbes into the PBS, then whole flower material was removed, and the tubes were centrifuged at 13k for 3 minutes. Top PBS was removed, and the remaining 100ul was used to resuspend the pelleted solids.

### Microbe isolation and identification

Brood cells, flowers, and water samples that were not used for DNA extraction were plated to isolate bacteria and fungi. These were plated either directly or after suspension in PBS onto both TSA +cycloheximide and YM+ chloramphenicol plates. Once colonies grew (1-5 days), representative colonies (based on morphology) were picked onto separate isolation plates of the same type and allowed to grow. Pure strains were then named based on isolation site (BH= Bodega Head, PR = Point Reyes (McClures); and saved as glycerol stocks in the −80. A subset of these were then identified via colony PCR (27F/1492R for bacteria, ITS1F/2 or NLR1/4 for fungi) followed by Sanger sequencing at the UC Davis DNA sequencing core, and NCBI BLAST.

### DNA Extraction

All samples were added whole to DNA extraction, following preprocessing. Extraction for all samples was done per manufacturer’s instructions with the DNeasy PowerSoil Pro kit. Four blanks were included in DNA extractions (Table 1). Extracted DNA was stored in the included extraction buffer at −80°C for amplicon sequencing and qPCR.

### Amplicon sequencing

Amplicon sequencing of extracted DNA was done to assess bacterial and fungal community composition using the 16S rRNA (V5/6) gene and ITS gene at the Integrated Microbiome Resource (IMR) at Dalhousie University in Halifax, Nova Scotia. Phusion Plus high-fidelity polymerase was used with fusion primers, which include the sequences below with Illumina adaptors + indices for multiplexing; sequencing was then performed on Illumina MiSeq (73, 74). Samples were de-multiplexed at IMR. For bacteria, primers 799F/1115R amplifying V5/V6 region of the 16S gene were used to limit mitochondria and chloroplast amplification (799F= 5’-AACMGGATTAGATACCCKG-3’/ 1115R= 5’-AGGGTTGCGCTCGTTG-3’)(75). These primers amplify a ∼300bp length target sequence. For fungi, primers ITS1F/ITS2 were used (ITS1F= 5’-CTTGGTCATTTAGAGGAAGTAA-3’/ ITS2=5’- GCTGCGTTCTTCATCGATGC-3’). These primers amplify the variable length ITS1/2 region.

### Amplicon data analysis

Sequences were analyzed in R (4.1.1)(76) with primarily the DADA2 package (1.22.0) (77), phyloseq (1.38.0) (78), vegan (2.6.4)(79), microbiome (1.23.1)(80) and ggplot2 (3.4.2)(81). See code for further details.

#### Bacteria

Reads were filtered and trimmed with the following parameters (others were default): maximum expected error was set to 2 for forward reads and 5 for reverse reads (to account for lower quality of reverse reads), reads were truncated at 280 and 220, respectively, to discard bases with quality scores <∼30). Primers were removed by trimming the respective primer length. Error rates, dereplication, denoising, merging, and chimera removal were done with default parameters; see supplemental code (Bacteria, code1) and data (‘16S_track_reads’). ASVs were inferred via DADA2 (1.22.0) and then taxonomy was assigned using the Silva 138.1 N99 database for bacteria (82). Mitochondria and chloroplast assigned reads were removed. Decontam package (1.14.0)(83) was used to identify and remove potential contaminants by comparing blanks to samples; 5 were found and removed at the threshold parameter of 0.5. Samples with less than 300 reads were then removed from further analysis, leaving N=86 samples; see supplemental code (Bacteria, code2). Samples lost (36) were: 4/4 blanks, 9/10 adult crop, 9/10 adult gut, 2/2 dirt (analyzed independently), 1/1 ice plant flower, 4/10 3rd-4^th^ instar larvae, 3/11 Summer prepupae, 4/10 Oct. prepupae, 1/7 pupae.

#### Fungi

Primers were removed using Cutadapt. Reads were filtered and trimmed using default parameters, aside from the length minimum, which was set to 70 to remove extremely short reads. Error rates, dereplication, denoising, merging, and chimera removal were done with default parameters; see supplemental code (Fungi, code1) and data (‘ITS_track_reads’). ASVs were inferred via DADA2 (1.22.0). We assigned fungal reads with the UNITE general release dynamic database (29.11.2022)(84). Non-fungal assigned reads were then removed, and Decontam package (1.14.0)(83) was used to remove potential contaminants by comparing blanks to samples; two were found and removed at threshold=0.5. Samples with fewer than 300 reads were then removed from further analysis, leaving N=93 samples. Samples lost (26) were: 4/4 blanks, 5/13 1st-2nd instar, 3/10 3rd-4th instar larvae, 1/1 dirt sample (analyzed independently), 9/10 adult crops, 1/10 adult guts, 1/1 gum flower, 2/7 pupae.

#### Community differences

To evaluate the compositional differences in microbial communities based on occurrence inside or outside of the brood cell, as well as for stage specific community separation, amplicon sequence data was used to create separate Bray-Curtis (BC) dissimilarity matrices for both bacteria and fungi. These were ordinated with NMDS (Fig. 2C,D). PERMANOVA was run based on BC distances to determine differences in community composition by sample type (in brood cell, out of brood cell, water, flowers) for both bacteria and fungi.

#### Relative abundance of Actinobacteria and *Moniliella* inside vs outside of brood cell

Data was subset to ASVs assigned to the Actinobacterial class (or *Moniliella spathulata* species). Samples were grouped based on whether they occur inside or outside of the brood cell (egg through unemerged adult: in brood cell; adult, flower and water samples: outside of brood cell). The total relative abundance of Actinobacteria (or *Moniliella spathulata*) assigned ASVs were compared between the groups with the Base R ‘stats’ package (4.1.1) (76) ‘kruskal.test’ function.

#### Defining the core microbiome

Detection of a core microbiome occurs using prevalence and abundance (detection) thresholds, and these can vary widely by study system, environment, and goals of the analysis (85, 86). Therefore, we used the ‘microbiome’ package “plot_core” function to visualize a wide range of prevalence (0 to 100%) and abundance (0.01% to 20%) thresholds for both bacteria and fungi in the style of a heatmap (Fig. 3) for clarity, and to allow for nuance in interpretation of what may be considered core taxa.

#### Abundance of *Streptomyces* by stage

Data was subset to ASVs assigned to the genus *Streptomyces*, and samples were grouped based on season, with egg-summer prepupa as ‘summer’, October and December prepupa as ‘overwintering’, and pupa-unemerged adults as ‘spring’. Differences in relative abundance of *Streptomyces* of these groups was evaluated with the Base R ‘stats’ package (4.1.1) (76) ‘kruskal.test’ function followed by the ‘FSA’ package (0.9.4)(87) ‘dunnTest’ function with Bonferroni p-value correction (88). To determine ‘actual’ abundance, we combined the qPCR data with the amplicon data by multiplying the total bacterial copy number by the proportion assigned to *Streptomyces* in each sample.

### qPCR- Microbial copy number

#### Bacteria

Bacterial copy number was quantified with standard DNA intercalating dye (SYBR) based qPCR. The same extracted samples that were sent for amplicon sequencing were run through this procedure. Identical primers (799F= 5’-AACMGGATTAGATACCCKG-3’ /1115R= 5’-AGGGTTGCGCTCGTTG-3’) were used so that compositional and quantification could be directly compared and merged. A 1:10 dilution of extracted DNA was determined after dilution testing was done with a representative subset of samples; 1:10 dilution gave in-range Cq values. Master mix, per reaction, was composed of 5ul SSO Advanced Universal SYBR Supermix (Catalog# 1725271), 0.3ul of each primer (10uM), 3.4ul Molecular grade water, and 1ul of extracted DNA (diluted 1:10 in Molecular grade water). Reactions were performed in triplicate for each sample, and arranged semi-randomly across plates to avoid possible correlations of plate and developmental stage. Blanks and standards were included in each plate, and a Cq cutoff for blanks was established at 31.

To translate Cq values to copy number, we purchased a plasmid containing the relevant sequence from *Nocardiodes luteus (*ASV_5) from Eurofins at a known concentration. This was diluted in 10-fold steps; the dilution steps of 1.17E+6 through 0.117 molecules/ul, plus a blank, were used to create a standard curve, which had an R^2^= 0.983. The equation of this line (see code) was used to convert Cq values to log (copy number), and edited to account for dilution.

Using amplicon data, we also adjusted the final qPCR copy number to remove the proportion of reads in each sample that had been assigned to mitochondria (no chloroplast reads were assigned). Flower and water samples had to be concentrated to ensure sufficient DNA for sequencing during pre-processing and thus bacteria in these samples could not be reliably quantified with qPCR.

#### Fungi

Fungal copy number was quantified with probe-based (Taq-Man®) qPCR using a previously established system, FungiQuant (33). Because the ITS region is highly variable in length, we did not use the same approach as above, as SYBR will intercalate throughout the length of an amplicon, potentially resulting in higher fluorescence readings for samples with a greater proportion of longer ITS amplicons. For these reasons, the fungal quantification was done with FungiQuant, using the 18S rRNA gene primers FungiQuant-F= 5′- GGRAAACTCACCAGGTCCAG-3′ and FungiQuant-R = 5′-GSWCTATCCCCAKCACGA-3′, along with the fluorescent probe FungiQuant-Prb = (6FAM) 5′-TGGTGCATGGCCGTT-3′ (MGBNFQ). As with bacteria, dilution testing of samples was done to bring Cq values into the optimal range, and a 1:20 dilution was picked. Master mix, per reaction, was composed of 5ul PCR Biosystems qPCRBIO Probe Mix (No-ROX) (Catalog# 17-512B), 0.3ul of each primer (10uM), 0.3ul fluorescent probe (10uM), 3.1ul molecular grade water, and 1ul of extracted DNA (diluted 1:20 in molecular grade water). Reactions were performed in triplicate for each sample, and arranged semi-randomly across plates to avoid possible correlations of plate and developmental stage. Blanks and standards were included in each plate.

To translate Cq values to copy number, we purchased a plasmid containing the relevant sequence from *Moniliella oedocephalis* on NCBI (#NG_062174.1) from Eurofins at a known concentration. This was diluted in 10-fold steps and the dilutions steps of 1.28E+6 through 0.128 molecules/ul, plus a blank, were used to create a standard curve, which had an R^2^= 0.983. The equation of this line (see code) was used to convert Cq values to log(copy number), and edited to account for dilution.

#### Statistical Analysis

To evaluate differences in copy number between stages for both bacteria and fungi, we used the Base R ‘stats’ package (4.1.1) (76) ‘kruskal.test’ function followed by the ‘FSA’ package (0.9.4)(87) ‘dunnTest’ function with BH p-value correction (89).

### Inhibition

#### Strain isolation

Strains of *Streptomyces* were isolated by plating brood cell contents on Tryptic Soy Agar (TSA) with added cycloheximide. Colonies were picked by hand and replated on TSA until isolated, then glycerol stocks were created. *Streptomyces* isolates were identified based on Sanger sequencing of the 16S rRNA gene (primers 27F/1492R) using NCBI BLAST (90).

*Thelonectria* was isolated in much the same way, but from an infected *A. bomboides* pupa that had developed external filamentous fungal growth. *Moniliella spathulata* was isolated from a 1^st^ instar *A. bomboides* provision (though it also occurred in nearly every plated brood cell sample, and all identified to same BLAST ID). *Ascosphaera apis* and *Aspergillus flavus* were isolated from infected *Bombus impatiens* larvae previously in the Vannette lab. All fungal isolation occurred with Yeast Media Agar with added chloramphenicol. Identification was based on Sanger sequencing of the ITS gene or lsrRNA D1/D2 region (primers ITS1F/ITS2 for *Thelonectria;* NL1/NL4 for *Ascosphaera apis*; ITS86F/ITS4 *Aspergillus flavus,* NL1/NL4 and ITS1/ITS4 for *Moniliella spathulata*) followed by NCBI BLAST (90) (Sequences in Table S1).

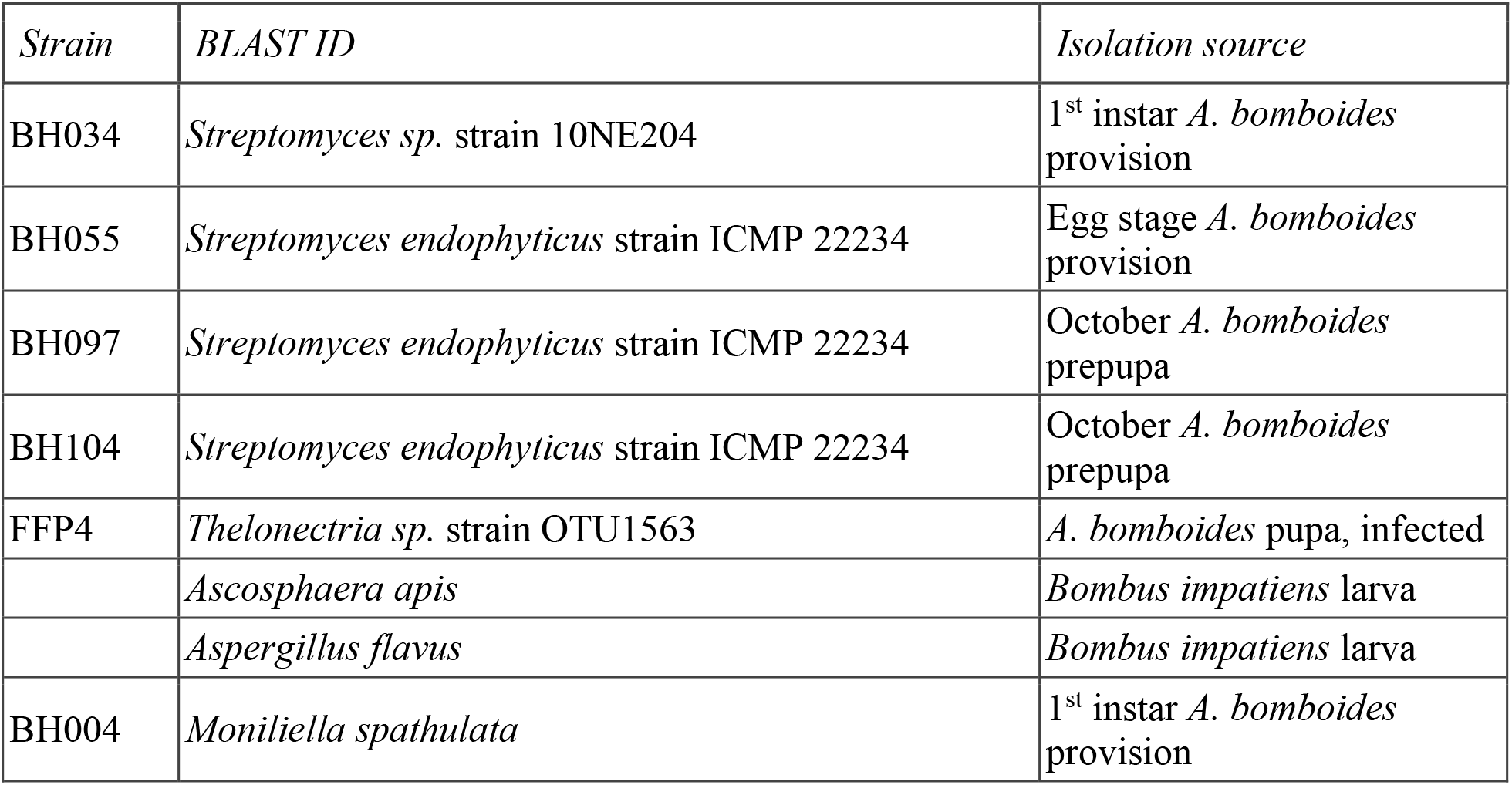

#### Trials

In order to ensure that both the bacteria and fungi could grow on the same media, for all inhibition trials we used TSA without any antimicrobials. For consistency, we used a template to mark the underside of all of the plates, it included in the very center a cross “+” then two parallel lines, 30mm long and each 20mm from the center point. These served as guides for inoculations. *Streptomyces* strains were inoculated with 1ul hoops from stock plates of the TSA without antifungals onto the 30mm parallel lines. Five replicate plates were made for each comparison (25 per trial, including 5 control plates). These were allowed to grow for 10 days, then, from stock plates of each fungus (also TSA, no antimicrobials), plugs were inserted into the center “+” of each plate. Care was taken to ensure that plugs were all taken from just inside the leading edge of the fungal hyphae on the stock plates. These were allowed to grow for seven days.

Measurements were taken on the backs of the plate, and measured the distance from the leading edge of the growing fungi to the center “+”, directly perpendicular to the parallel lines, on both sides. A tabletop light pad was used for imaging to qualitatively assess the density of the fungal hyphae, ensuring even back-lighting for the plates.

#### Analysis

Radius measurements (two per plate, each side of the ‘+’) were averaged for each replicate plate. Kruskal-Wallis was run with the Base R ‘stats’ (4.1.1) package ‘kruskal.wallis’ function as radius by inhibition treatment. This was followed with multiple comparisons with the ‘FSA’ package (0.9.4)(87) ‘dunnTest’ function, but as we were only interested in comparisons to the negative control, we then subset to those four comparisons. P value correction done with ‘stats’ package ‘p.adjust’ function using a Bonferroni correction (88).

### Sugar and Sugar Alcohols

Samples of whole larvae, prepupae and pupae, as well as one pollen provision from a 4^th^ instar larva were extracted for sugar and sugar alcohol analysis. Whole samples were placed in tubes with metal beads and 1mL of 100% ethanol and run on a bead beater for 8 minutes at full speed with 20s breaks every minute. These were then centrifuged for 30 seconds at 10k rcf. For each sample, the top 700ul of ethanol was moved to a new tube, 700ul 100% hexane was added, and then vortexed for 30 seconds. To this, 100ul MilliQ water was added, and vortexed for another 30 seconds. Once hexane had separated from the aqueous phase, it was removed (800ul). The remaining 1mL of aqueous phase was centrifuged for 2 minutes at 16k rcf, and the bottom 500ul was filtered through a 0.2 micron syringe filter and placed in a new tube in a lyophilizer for 6 hours, without heat. The dried samples were kept in a −20C freezer until analysis, at which time they were re-suspended in 300ul 1:1 water: acetonitrile. Standards of erythritol, sorbitol, fructose, glucose, sucrose, xylose and maltose were made at 0.5 mg/mL, standards of glycerol and trehalose were made at 5mg/mL and 1mg/mL respectively, all in 1:1 water: acetonitrile.

Separation of sugars was performed on Thermo UltiMate 3000 HPLC system according to the Waters Application Note: WA60110, except for the following: column was Phenomenex Luna Omega 3um SUGAR (50×2.1mm, Part#: 00B-4775-AN), and flow rate was 0.2mL/min; detection was by CAD (Corona Veo; Dionex). Each sample was run twice, standards were run 2-5 times. Analysis of peaks was performed with Thermo Fisher Chromeleon software. Peak identities were assigned based on retention times of standards, and unassigned peaks were then named by their retention times. Peak area was calculated by the software and this data was exported for analysis.

To identify differences in sample groups based on SSA profiles we used Principal Components Analysis (PCA) ‘stats’ package, Base R (76). Data was first normalized by Hellinger transformation. The ‘factoextra’ package was used to plot PCA and biplot of components. After calculation of Bray-Curtis distance matrix, PERMANOVA (‘vegan’ package; (79) and pairwise PERMANOVA (package ‘RVAideMemoire’ 0.9.83) were used to determine differences in composition of SSA by sample group, p-value correction by FDR (89).

## Author contributions

SMC, QSM, BD, SLB, and RLV contributed to selection of this bee and floral study system, sampling design and fieldwork, SC performed lab work with contributions by SS, SC performed bioinformatic and statistical analysis, SC and RLV wrote the paper and all authors contributed to revisions.

