## Supplemental Figures and Tables for "Microbial Metamorphosis: Symbiotic bacteria and fungi proliferate during diapause and may enhance overwintering survival in a solitary bee"

Christensen *et al* 2023

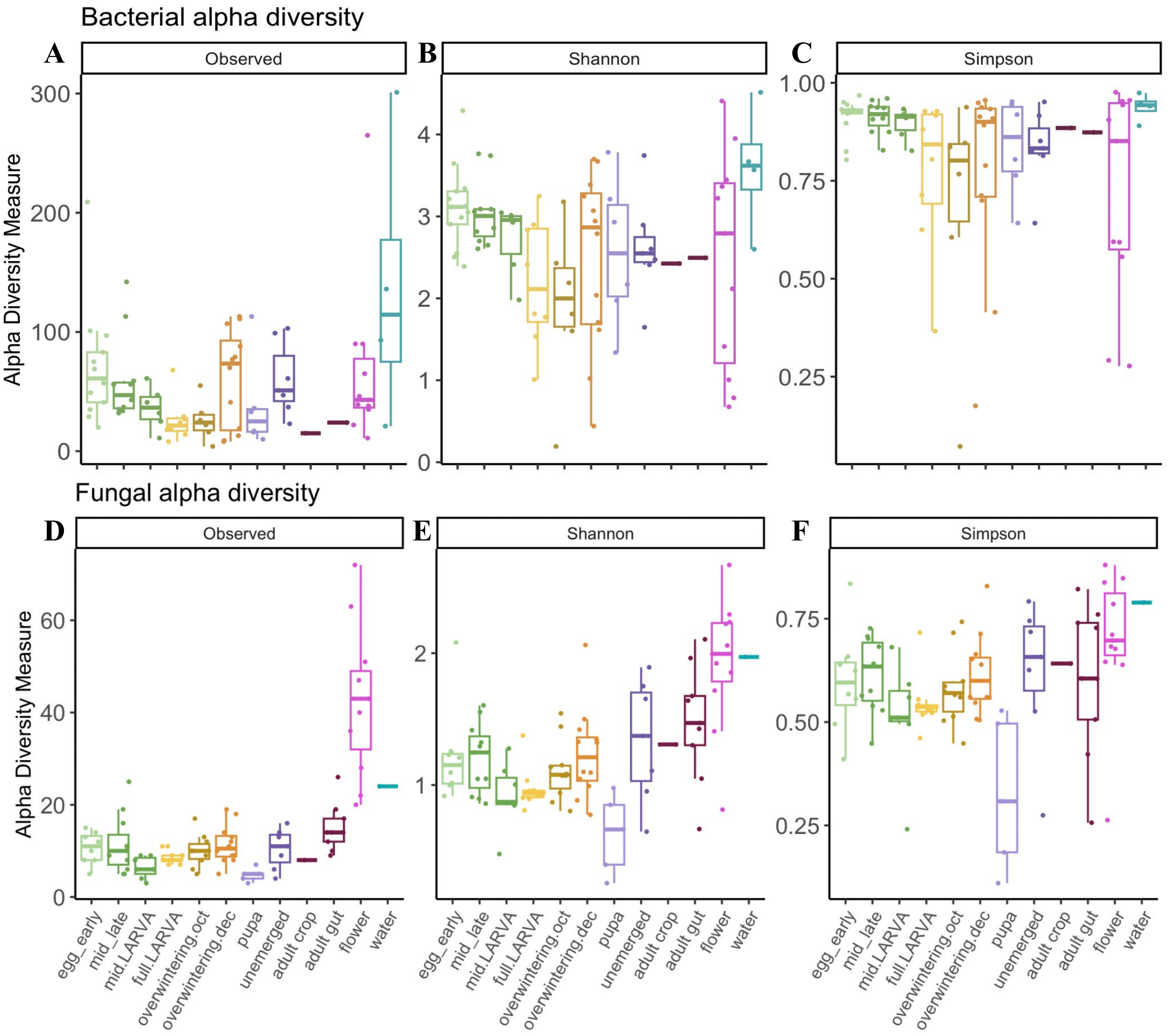


**Fig. S1- Bacterial and fungal alpha diversity by developmental stage and sample type.** Alpha diversity metrics (y axis) by sample type (x axis). No ASVs removed. **(A-C)** Bacterial alpha diversity: Observed alpha diversity sample type (x axis) is significant globally but no comparisons remain significant after p-value correction. (Kruskal-Wallis $X^2$= 22.838, df = 11, p= 0.01863). Shannon diversity metric does not differ globally by sample type (Kruskal-Wallis $X^2$= 16.616, df = 11, p-value > 0.05). Simpson diversity metric does not differ globally by sample type (Kruskal-Wallis $X^2$ = 15.295, df = 11, p-value > 0.05). **(D-F)** Fungal alpha diversity varies significantly by sample type in all measurements (Observed: Kruskal-Wallis $X^2$= 51.8, df = 11, p-value = 3e-7; Shannon: Kruskal-Wallis $X^2$= 39.4, df = 11, p-value =4.5e-5; Simpson: Kruskal-Wallis $X^2$ = 26.9, df = 11, p-value = 0.005).

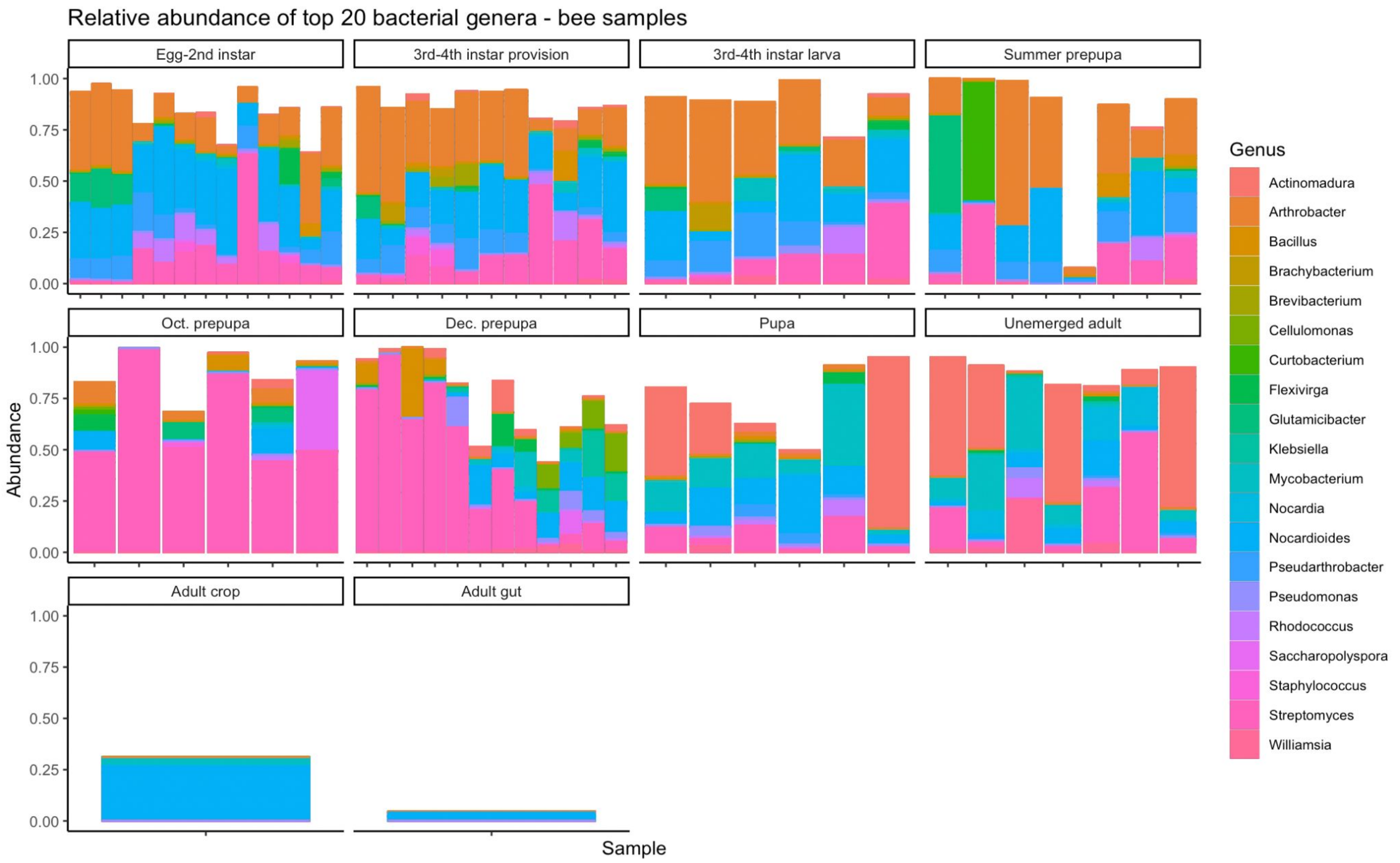


**Fig. S2- Top 20 bacterial genera comprise the majority of reads in each stage, in varying abunances.** Data subset to include only ASVs belonging to the 20 most abundant genera, and the white space indicates proportion of sample comprised by additional genera. ASVs are grouped and colored by genus, shown as relative abundance in sample. Each bar represents one sample, and samples are grouped by stage.

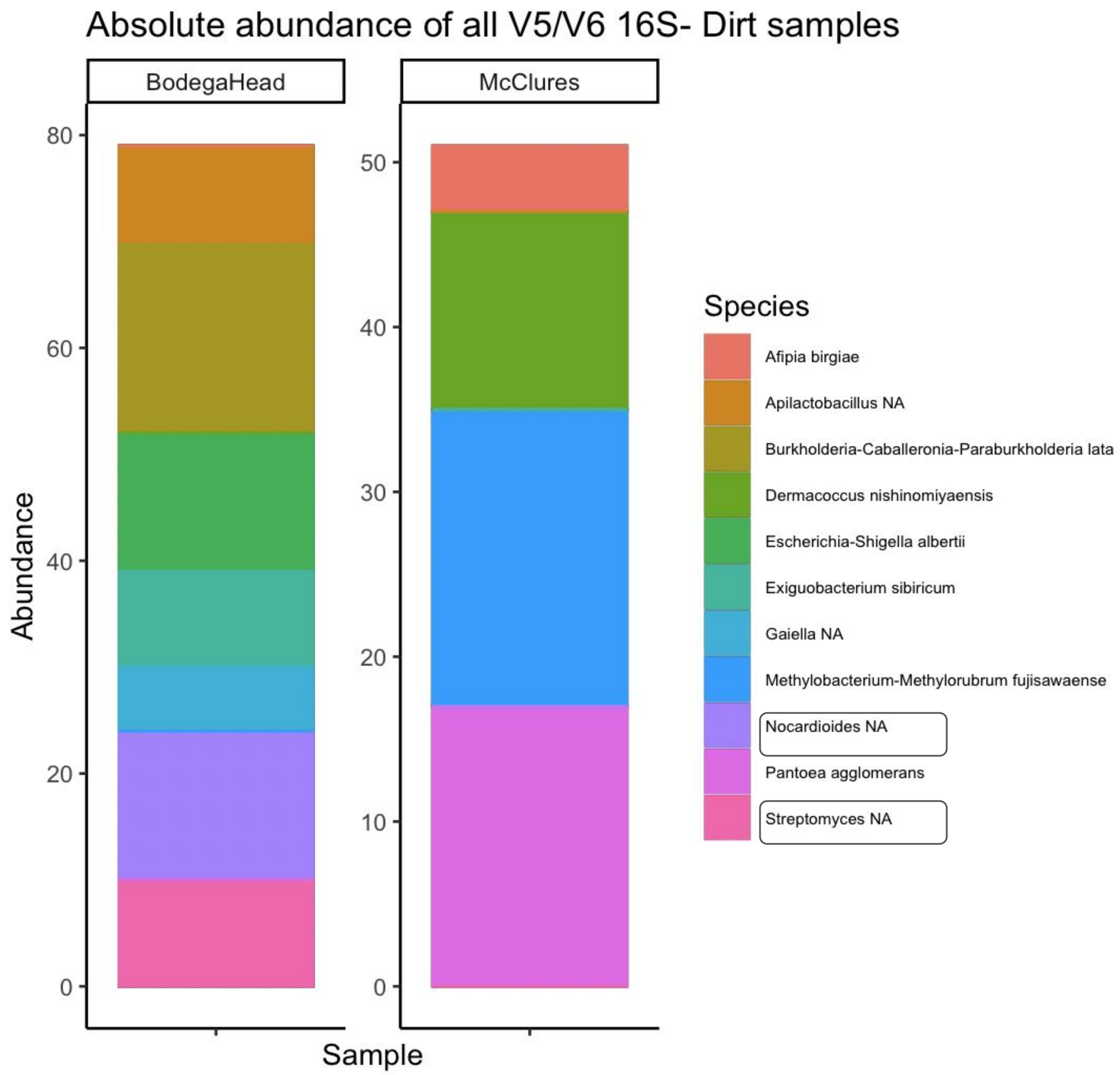


**Fig. S3- Soil sample microbial composition.** Soil samples were collected from each site, from ~2" beneath the surface, within the area of the nesting aggregation but not touching any nest. They were analyzed separately from the rest of the samples due to very low read count (note y axis), using a less stringent pipeline, and without using Decontam. *Nocardiodes* and *Streptomyces* (circled in key) were present in soil at Bodega Head but not found at McClure's. In separate analysis of fungal reads, no reads passed merge step: manual analysis showed there were 20 total reads in the McClure's soil sample, two of these reads were *M. spathulata*, the rest were non-fungi.

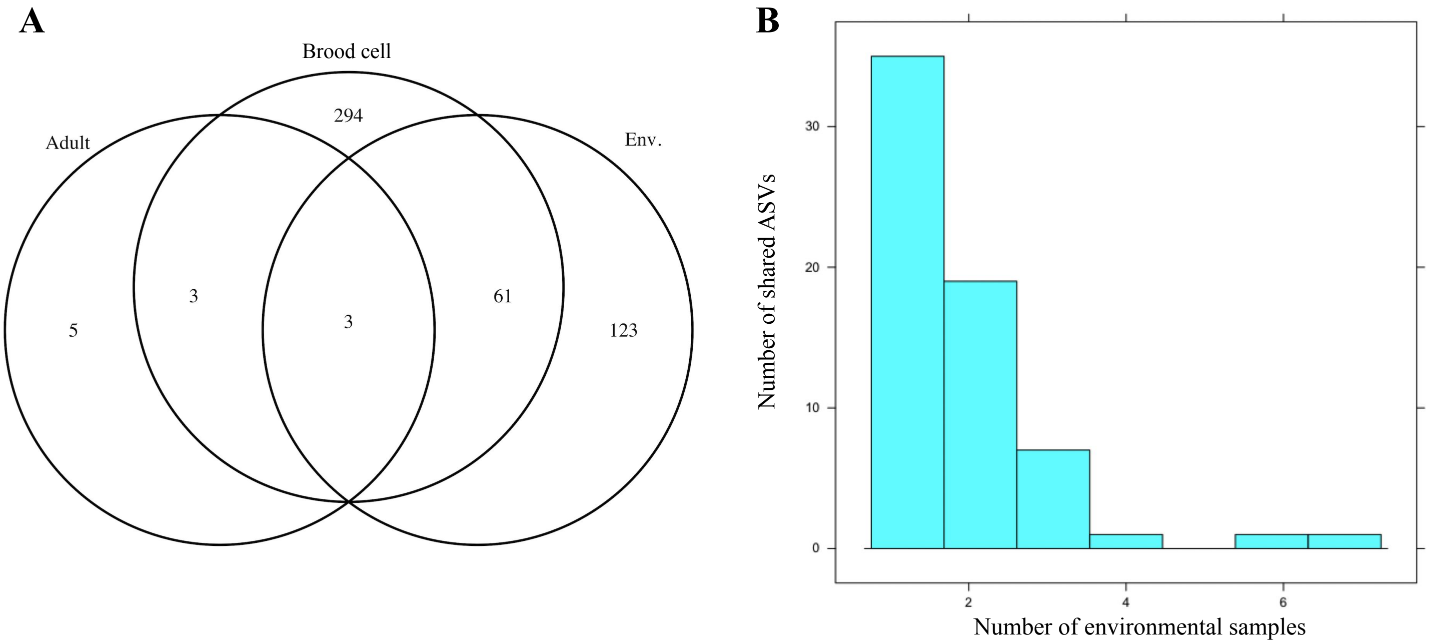


**Fig. S4- Overlap of Actinobacterial ASVs with environmental ASVs does not fully explain acquisition.** To assign ASV presence within a sample, we used a cutoff of 0.1% relative abundance, and filtered to only ASVs assigned to Actinobacteria.

**A)** Three ASVs: ASV_29, ASV_78, ASV_471 (Assigned as *Nocardioides*, *Mycobacterium*, and order: Frankiales, respectively) were found to occur in all sample groups (eg, adult, brood cell, environment). ASV_29 was found in 53.2% of all samples, ASV_78 was found in 23.9% of all samples, ASV_471 found in 4.3% of all samples. The mean relative abundances (in samples where the ASV was found) are ASV_29 mean= 0.017 SD= 0.014; ASV_78 mean = 0.015, SD= 0.015; ASV471 mean= 0.0013 SD=0.01. In total, 64 Actinobacteria ASVs overlap between brood cell and environmental samples, which represents less than one fifth (17.7%) of the Actinobacterial ASVs found in brood cell samples. Brood cell n=69, environment n=15, adult n=2.

**B)** Histogram of the 64 ASVs shared between brood and environmental samples, showing their representation in environmental samples. The vast majority (54 ASVs, 90%) of these ASVs were found in only one or two environmental samples (the first two bars on the left). The environmental samples contributing the most shared Actinobacterial ASVs were: Radish flower (15x flowers bulked) which contained 31 shared ASVs, and Sea Daisy (3x flowers bulked), which contained 27 shared ASVs.

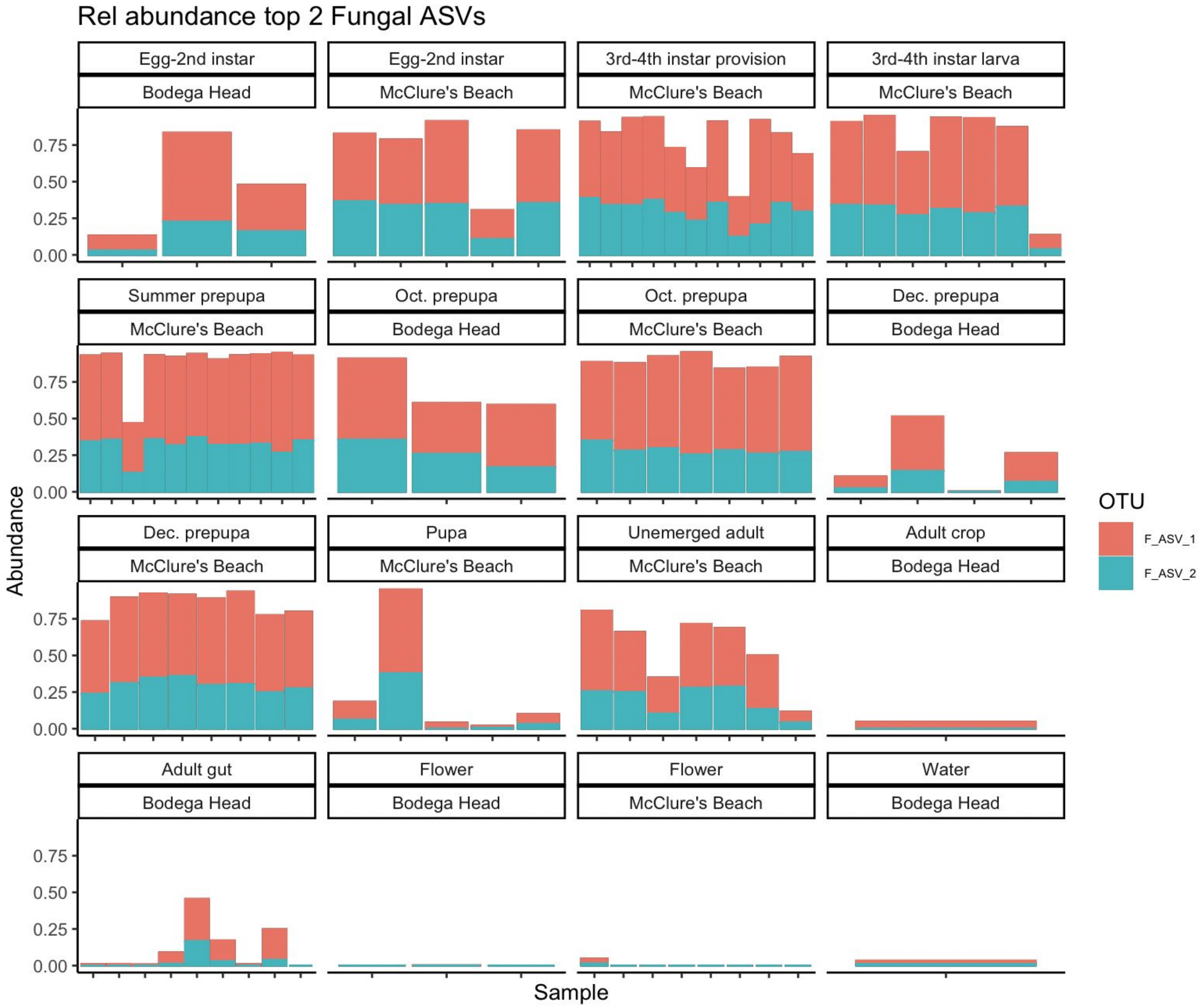


**Fig. S5- Two ASVs of *Moniliella spathulata* comprise the majority of reads in most brood cell samples and are found at both sites.** Relative abundance data is shown, subset only to the top 2 ASVs in the fungal dataset, which are both assigned to *Moniliella spathulata*. Each bar represents one sample, samples are separated by stage or sample type and by collection site.

| **Table S1- Sequences of bacterial and fungal isolates used in inhibition assays** | | |
|---|---|---|
| *Strain* | *BLAST ID & Sequence* | *Isolation source* |
| BH034 | *Streptomyces sp.* strain 10NE204 | 1st instar *A. bomboides* provision |
| | 27F/1492R:5’-GGGACAAGCCTTGGAAACGAGGTCTAATACCGGATAACACTCCCATTCTCCTGGATGGGGGTTAAAA<br>GCTCCGGCGGTGAAGGATGAGCCCGCGGCCTATCAGCTTGTTGGTGAGGTAATGGCTCACCAAGGCGACGACGGG<br>TAGCCGGCCTGAGAGGGCGACCGGCCACACTGGGACTGAGACACGGCCCAGACTCCTACGGGAGGCAGCAGTGGG<br>GAATATTGCACAATGGGCGAAAGCCTGATGCAGCGACGCCGCGTGAGGGATGACGGCCTTCGGGTTGTAAACCTCT<br>TTCAGCAGGGAAGAAGCGMAAGTGACGGTACCTGCAGAAGAAGCGCCGGCTAACTACGTGCCAGCAGCCGCGGT<br>AATACGTAGGGCGCAAGCGTTGTCCGGAATTATTGGGCGTAAAGAGCTCGTAGGCGGCTTGTCACGTCGATTGTGA<br>AAGCTCGGGGCTTAACCCCGAGTCTGCAGTCGATACGGGCTAGCTAGAGTGTGGTAGGGGAGATCGGAATTCCTGG<br>TGTAGCGGTGAAATGCGCAGATATCAGGAGGAACACCGGTGGCGAAGGCGGATCTCTGGGCCATTACTGACGCT<br>GAGGAGCGAAAGCGTGGGGGAGCGAACAGG-3’ | |
| BH055 | *Streptomyces endophyticus* strain ICMP 22234 | Egg stage *A. bomboides* provision |
| | 27F/1492R:5’-AGTCGAACGATGAAGCCCTTCGGGGTGGATTAGTGGCGAACGGGTGAGTAACACGTGGGCAATCTGC<br>CCTTCACTCTGGGACAAGCCTTGGAAACGAGGTCTAATACCGGATAACACTCCCATTCTCCTGGATGGGGGTTAAA<br>AGCTCCGGCGGTGAAGGATGAGCCCGCGGCCTATCAGCTTGTTGGTGAGGTAATGGCTCACCAAGGCGACGACGG<br>GTAGCCGGCCTGAGAGGGCGACCGGCCACACTGGGACTGAGACACGGCCCAGACTCCTACGGGAGGCAGCAGTGG<br>GGAATATTGCACAATGGGCGAAAGCCTGATGCAGCGACGCCGCGTGAGGGATGACGGCCTTCGGGTTGTAAACCT<br>CTTTCAGCAGGGAAGAAGCGAAAGTGACGGTACCTGCAGAAGAAGCGCCGGCTAACTACGTGCCAGCAGCCGCGG<br>TAATACGTAGGGCGCAAGCGTTGTCCGGAATTATTGGGCGTAAAGAGCTCGTAGGCGGCTTGTCACGTCGATTGTG<br>AAAGCTCGGGGCTTAACCCCGAGTCTGCAGTCGATACGGGCTAGCTAGAGTGTGGTAGGGGAGATCGGAATTCCTG<br>GTGTAGCGGTGAAATGCGCAGATATCAGGAGGAACACCGGTGGCGAAGGCGGATCTCTGGGCCATTACTGAC-3’ | |
| BH097 | *Streptomyces endophyticus* strain ICMP 22234 | October *A. bomboides* prepupa |
| | 27F/1492R:5’-GGGTGGATTAGTGGCGAACGGGTGAGTAACACGTGGGCAATCTGCCCTTCACTCTGGGACAAGCCTT<br>GGAAACGAGGTCTAATACCGGATAACACTCCCATTCTCCTGGATGGGGGTTAAAAGCTCCGGCGGTGAAGGATGA<br>GCCCGCGGCCTATCAGCTTGTTGGTGAGGTAATGGCTCACCAAGGCGACGACGGGTAGCCGGCCTGAGAGGGCGA<br>CCGGCCACACTGGGACTGAGACACGGCCCAGACTCCTACGGGAGGCAGCAGTGGGGAATATTGCACAATGGGCGA<br>AAGCCTGATGCAGCGACGCCGCGTGAGGGATGACGGCCTTCGGGTTGTAAACCTCTTTCAGCAGGGAAGAAGCGA<br>AAGTGACGGTACCTGCAGAAGAAGCGCCGGCTAACTACGTGCCAGCAGCCGCGGTAATACGTAGGGCGCAAGCGT<br>TGTCCGGAATTATTGGGCGTAAAGAGCTCGTAGGCGGCTTGTCACGTCGATTGTGAAAGCTCGGGGCTTAACCCCG<br>AGTCTGCAGTCGATACGGGCTAGCTAGAGTGTGGTAGGGGAGATCGGAATTCCTGGTGTAGCGGTGAAATGCGCA<br>GATATCAGGAGGAACACCGGTGGCGAAGGCGGATCTCTGGGCCATTACTGACGCTGAGGAGCGAAA-3’ | |
| BH104 | *Streptomyces endophyticus* strain ICMP 22234 | October *A. bomboides* prepupa |
| | 27F/1492R:5’-GGATTAGTGGCGAACGGGTGAGTAACACGTGGGCAATCTGCCCTTCACTCTGGGACAAGCCTTGGAA<br>ACGAGGTCTAATACCGGATAACACTCCCATTCTCCTGGATGGGGGTTAAAAGCTCCGGCGGTGAAGGATGAGCCCG<br>CGGCCTATCAGCTTGTTGGTGAGGTAATGGCTCACCAAGGCGACGACGGGTAGCCGGCCTGAGAGGGCGACCGGC<br>CACACTGGGACTGAGACACGGCCCAGACTCCTACGGGAGGCAGCAGTGGGGAATATTGCACAATGGGCGAAAGCC<br>TGATGCAGCGACGCCGCGTGAGGGATGACGGCCTTCGGGTTGTAAACCTCTTTCAGCAGGGAAGAAGCGAAAGTG<br>ACGGTACCTGCAGAAGAAGCGCCGGCTAACTACGTGCCAGCAGCCGCGGTAATACGTAGGGCGCAAGCGTTGTCC<br>GGAATTATTGGGCGTAAAGAGCTCGTAGGCGGCTTGTCACGTCGATTGTGAAAGCTCGGGGCTTAACCCCGAGTCT<br>GCAGTCGATACGGGCTAGCTAGAGTGTGGTAGGGGAGATCGGAATTCCTGGTGTAGCGGTGAAATGCGCAGATAT<br>CAGGAGGAACACCGGTGGCGAAGGCGGATCTCTGGGCCATTACTGACGCTGAGGAGCGAAAGCGTGGGGAGCGA<br>A-3’ | |

**Table S1 (continued)**

| *Strain* | *BLAST ID & Sequence* | *Isolation source* |
|---|---|---|
| FFP4 | *Thelonectria sp.* strain OTU1563 | *A. bomboides* pupa, infected |
| | ITS1F/ITS2:5'-CCCCGCCAGTACTCTGGCGGCATGCCTGTTCGAGCGTCATTACAACCCTCAGCCCCCGGGCTTGGCG TTGGGGATCGGCACAAGGCGGAGGGGCAMCCCTAGGGAGCCCCCTCCGGCCCGCCGTCCCCCAAATTCAGTGGCG GTCACCGTCGCGGCCTCCTATGCGTAGTAGCAACACCTCGCACTGGAARMMCCCTCGSGTCCACGCCGTAAAACCC CCGACTTTTTTTATCAAGTTGACCCTCGAA-3' | |
| *A.apis* | *Ascosphaera apis* | *Bombus impatiens* larva |
| | NL1/NL4:5'-ATCCGTGCCGAGCGCGTTCCTCAGTCCCGGCTGGCCGCATTGCACCTCGGGCTATAAGACCTCCCGGA GGAGGATACATTCCCGAAGCCTTTAACCGGCCGCCAGAACTGATGCTGGCCCGACCGCAGGGGAATACACAGGGG AGAACCCCTGCTGAACCCCCACGGCCGACTCTGGTCGCATGTGCTTCCCTTTCAACAATTTCACATGCTTTTTAACT CTCTTTTCAAAGTGCTTTTCATCTTTCGATCACTCTACTTGTGCGCTATCGGTCTCCGACCAGTATTTAGCTTTAGAT GAAATTTACCACCCATTTAGAGCTGCATTCCCAAACAACTCGACTCGTCGAAGGAGCTCTACACGGTTGCAGCCGG CCGGCCGAAGACGGGATTCTCACCCTCTGCGACGCCCCGTTCCAGGGGACTTAGACCGCGACCACTCCCAGAGCAT CCTCTCCAAATTACAACTCGGACCCCGAAGGAGCCAGATTTCAAATTTGAGCTTTTGCCGCTTCACTCGCCGTTACT GAGGCAATCCCTGTTGGTTTCTTT-3' | |
| *A. flavus* | *Aspergillus flavus* | *Bombus impatiens* larva |
| | ITS86F/ITS4:5'-ATGCCTGTCCGAGCGTCATTGCTGCCCATCAAGCACGGCTTGTGTGTTGGGTCGTCGTCCCCTCTCC GGGGGGGACGGGCCCCAAAGGCAGCGGCGGCACCGCGTCCGATCCTCGAGCGTATGGGGCTTTGTCACCCGCTCT GTAGGCCCGGCCGGCGCTTGCCGAACGCAAATCAATCTTTTTCCAGGTTGACCTCGGATCAGGTAGGGATACCCGC TGAACTTAAGCATATCAAAAAGCGGAGG-3' | |
| BH004 | *Moniliella spathulata* | 1st instar *A. bomboides* provision |
| | ITS1F/ITS2:5'-TCCTGGTTATTTCTCACCTGAGGGTTCGCCCTCTCTCTCACACTGTGAACAAAACAAAACGAAATTGA GGATGGGAAGTATTTTTTAGTAAAAAAAACAACTTTTGGCAATGGATCTCTTGGTTCTCCCATCGATGAAGAACGC AGCGAATCGCGAAAGGTAGTGTGAATTGCAGAACCGTGAATCATCGAGTCTTTGAACGCATCTTGCGCCAGGGGGC CATTCCCCCCGGCATGCCTGTTTGAGTGTTGTGAATCTCTACCTAGCCGCGCCAGGCTAGGATGTGGGTTTGCTCCT TCTCGGAGCTCACCTGAAAAGAAAAAGGGCACACTGGATATGGTGACACATTGTATCTCAGAAACGCCTCTCTTAC ACTTTCGACCTCAGATCAGGTAGGACTACCCGCTGAACTTAAGCATA-3'<br>NL1/NL4:5'-AGCGAAGCGGGATGAGCCCAGATTTGAAAGCTCCCGCCAGGGCGCATTGTAAGCTGGAGACGTGCCT CGAGCGGCGCGGCTGGACGCAAGTCTGCTGGAAAGCAGCATCAGAGAGGGTGAGAATCCCGTGCTTGGTCTGGCT GTGCGCCGTGTTGTGGGGTGCGCTCGACGAGTCGCGTTGTTTGGGAATGCAGCGCAAAGAGGGGTGGTAAACGCC ATCCAAGGCTAAATACCGGGGAGAGACCGATAGCGAACAAGTACCGTGAGGGAAAGATGAAAAGCACTTTGGAA AGAGAGTTAAAGAGTACGTGAAATTGCCAAGAGGGAAGCGCTGGCAGTCAGTGCCGTAGCGCTGCTGGTCCCGCC TTTTTTTGGGAGGTTGATGCCGGCAGTGTGGGGCCCGCGTCGGTTGCTGTTTTGGTGGGGGAGAAGGCAGAGCGGA AGGTGGCTTCCCTTTTTTTGGGGGGAGTGTTATAGCCGCTTTGTGGATGCCCTGCTAGCGACCGAGGACCGCTTTTT TCGATAGAGGATGCGGGCTTAATGGCTTT-3' | |
